# Uridine Phosphorylase-1 supports metastasis of mammary cancer by altering immune and extracellular matrix landscapes of the lung

**DOI:** 10.1101/2024.07.02.601676

**Authors:** Declan Whyte, Johan Vande Voorde, David Sumpton, Sandeep Dhayade, Emmanuel Dornier, Madeleine Moore, David Novo, Jasmine Peters, Robert Wiesheu, John B.G. Mackey, Amanda J. McFarlane, Frédéric Fercoq, Sophie Fisher, Carolina Dehesa Caballero, Kathryn Gilroy, Keara L. Redmond, Louise E. Mitchell, Eve Anderson, Gemma Thomson, Lindsey N. Dzierozynski, Juan J. Apiz Saab, Caroline A. Lewis, Alexander Muir, Christopher J. Halbrook, Douglas Strathdee, Rene Jackstadt, Colin Nixon, Philip Dunne, Colin W. Steele, Leo M. Carlin, Iain R. Macpherson, Edward W. Roberts, Seth B. Coffelt, Karen Blyth, Owen J. Sansom, Jim C. Norman, Cassie J. Clarke

**Affiliations:** Cancer Research UK Scotland Institute, Garscube Estate, Switchback Road, Glasgow, G61 1BD, UK; School of Cancer Sciences, University of Glasgow, Garscube Estate, Switchback Road, Glasgow, G61 1QH, UK; Queen’s University Belfast, University Rd, Belfast, BT7 1NN, UK; Ben May Department for Cancer Research, University of Chicago, Chicago IL USA; Whitehead Institute for Biomedical Research, Cambridge, MA, USA; Department of Molecular Biology and Biochemistry, University of California Irvine, Irvine, CA, USA; Chao Family Comprehensive Cancer Center, University of California Irvine, Orange, CA, USA

**Keywords:** Breast Cancer, Neutrophils, Lung Metastasis, Pyrimidine Metabolism, Uridine Phosphorylase-1, Uracil, T cells, Fibronectin

## Abstract

Understanding the mechanisms that facilitate early events in metastatic seeding is key to developing therapeutic approaches to reduce metastasis – the leading cause of cancer-related death. Using whole animal screens in genetically engineered mouse models of cancer we have identified circulating metabolites associated with metastasis. Specifically, we highlight the pyrimidine uracil as a prominent metastasis-associated metabolite. Uracil is generated by neutrophils expressing the enzyme uridine phosphorylase-1 (UPP1), and neutrophil specific *Upp1* expression is increased in cancer. Altered UPP1 activity influences expression of adhesion molecules on the surface of neutrophils, leading to decreased neutrophil motility in the pre-metastatic lung. Furthermore, we find that UPP1-expressing neutrophils suppress T-cell proliferation, and the UPP1 product uracil can increase fibronectin deposition in the extracellular microenvironment. Consistently, knockout or inhibition of UPP1 in mice with mammary tumours increases the number of T-cells and reduces fibronectin content in the lung and decreases the proportion of mice that develop lung metastasis. These data indicate that UPP1 influences neutrophil behaviour and extracellular matrix deposition in the lung and suggest that pharmacological targeting of this pathway could be an effective strategy to reduce metastasis.

## Introduction

Metastasis is the process whereby cancer cells move from their original location in the primary tumour to distant sites in secondary organs. It is responsible for >90% of cancer-related deaths [1, 2] and thus represents a pressing area of unmet clinical need. Whilst metastatic disease may be detected at original diagnosis, more often it occurs as recurrent disease downstream of primary presentation. This is a significant problem for patients diagnosed with breast cancer, a heterogenous disease of different subtypes where relapse can occur across wide-ranging timescales from months to decades after primary tumour surgery [3]. Understanding mechanisms that facilitate metastasis is therefore important, not only to identify new targets for therapeutic treatment strategies, but to also identify the patients at highest risk of developing metastatic disease, ensuring high-risk patients are stratified for appropriate treatment regimens to reduce their risk of recurrence, whilst those with indolent disease may be spared aggressive over-treatment.

Metastasis occurs through a series of well-defined steps [4], culminating in outgrowth of cancer cells in distant organs. Factors released into the circulation by primary tumours can evoke alterations at metastatic target sites prior to the arrival of disseminated cancer cells, leading to formation of pre-metastatic niches which facilitate metastatic seeding. Whilst the immune system can contribute to tumour surveillance and elimination of incipient cancers, primary tumours can also evoke inflammatory responses which promote tumour growth and invasiveness. Furthermore, tumour-induced alterations in immune landscapes also contribute to formation of pre-metastatic niches. For example, primary tumour-driven mobilisation of neutrophils from the bone marrow to distant target sites promotes vascular leakiness which supports extravasation [5, 6], whilst increased numbers of neutrophils in metastatic target organs creates immunosuppressed microenvironments that enable disseminated cancer cells to evade immune challenge, thus facilitating metastatic outgrowth [7–12]. Neutrophils also play key roles in mediating cell-cell communication within pre-metastatic niches, attracting additional cell types and supplying endopeptidases to facilitate remodelling of the extracellular matrix (ECM) [13]. Indeed, structural changes in the composition, organisation and mechanical properties of the ECM can further contribute to establishment of microenvironments that enable cancer cells to thrive [14].

Altered metabolism can also influence metastasis, and whilst a predominant focus has been on the role of cancer cell-intrinsic metabolism in this process [15], the nature of the metastatic cascade means that the metabolism of other cell types in local and distant microenvironments may also be important. Resident mesenchymal cells in the pre-metastatic lung have been shown to promote metastasis of breast cancer through metabolic reprogramming of tumour cells and natural killer (NK) cells [16], whilst neutrophils, which are traditionally thought of as being highly glycolytic [17], have been shown to engage mitochondrial metabolism to circumvent nutrient limitations and maintain immune suppression [18]. As a consequence, altered immune landscapes [19], ECM deposition [20] and dysregulated metabolism [21] have all been associated with poor prognosis for cancer patients, and whilst these mechanisms may have the ability to influence disease independently, it is likely that they will also conspire to collectively facilitate metastatic progression.

In characterising serum metabolites from genetically engineered mouse models (GEMMs) of metastatic cancer, we find that hallmarks of the metastasis-associated circulating metabolome can be generated by mobilisation of pro-metastatic neutrophils. Specifically, we show that a key aspect of pyrimidine metabolism (uridine phosphorylase-1 (UPP1)-catalysed phosphorolysis of uridine into uracil and ribose-1 phosphate) influences priming of the lung metastatic niche, and consequent metastasis of mammary cancer to this organ. Mechanistically, our data highlight novel roles for UPP1 in influencing immune landscapes and ECM deposition in the metastatic lung. Importantly, we find that the ability of mammary cancer to metastasise to the lungs is reduced in the absence of UPP1. Our data, therefore, highlight a previously unknown and unexpected role for UPP1 in facilitating metastasis.

## Results

### Circulating uracil correlates with metastasis in mouse models of cancer

To characterise the circulating metabolome of metastatic cancer we performed liquid chromatography-mass spectrometry (LC-MS) on polar metabolites extracted from the serum of cancer GEMMs. To model metastasis of mammary cancer we used the MMTV-*PyMT* mouse model, in which polyomavirus middle T antigen is expressed specifically in the mammary epithelium resulting in formation of mammary adenocarcinomas that recapitulate the progression of human breast cancer subtypes with poor prognoses [22–24]. Importantly, when expressed in an FVB/N mouse strain, this results in mammary tumours with high propensity to metastasise to the lungs [25]. Comparing serum from MMTV-*PyMT* female mice bearing mammary tumours to age-matched non-tumour bearing FVB/N controls, 214 compounds were identified as being significantly altered in mammary tumour bearing animals (Figure S1). The extent of metastatic burden in the lungs of tumour-bearing mice was quantified, and a signature of circulating metabolites which correlated with metastasis (but not primary tumour burden) was identified (Figure 1A; Figure S1B). This indicated that 17 compounds had positive, and 19 compounds had negative, correlations with metastatic burden independent from any correlation with primary tumour burden (Table S1). Parallel metabolic characterisation was also performed in GEMMs of other cancer types. For colorectal cancer, KPN mice were used (*villin*Cre^ER^; *Kras*^G12D/+^; *Trp53*^fl/fl^; *R26*^N1icd/+^), a model in which NOTCH1 activation is induced specifically in the intestinal epithelium to generate highly metastatic *Kras^G12D^*-driven cancer; equivalent to the serrated colorectal cancer subtype [10]. KPN mice at clinical endpoint (100% metastatic penetrance) were compared with KPN mice 30 days post induction (no metastatic disease). For pancreatic adenocarcinoma (PDAC), KPC mice (*Pdx1*-Cre; LSL-*Kras*^G12D/+^; LSL-*Trp53*^R172H/+^) were compared to KP^fl^C mice (*Pdx1*-Cre; LSL-*Kras*^G12D/+^; LSL-*Trp53*^fl/+^). In these models, pancreas-specific expression of *Kras*^G12D^ leads to the development of pancreatic tumours with similar penetrance across both mouse cohorts, however those with mutant p53 (KPC) develop highly metastatic disease, whereas those with p53 loss (KP^fl^C) develop PDAC that has a tendency not to metastasise [26]. For KPC and KP^fl^C mice, comparisons were made at palpable primary tumour, a timepoint in which pro-metastatic structural changes are apparent in distant target organs but that precede the formation of overt metastases [27], providing the opportunity to identify metabolites that may contribute to metastatic niche priming. Assessment of serum LC-MS data generated from all these GEMMs identified uracil as a metabolite that was consistently detected in metastatic mammary cancer (Figure 1B-C) and colon cancer (Figure 1D), and metastatic niche priming of PDAC (Figure 1E). Furthermore, analysis of plasma from patients with metastatic breast cancer indicated that increased levels of circulating uracil were present in the circulation of breast cancer patients with metastatic disease by comparison with healthy volunteers (Figure 1F).

**Figure 1:**
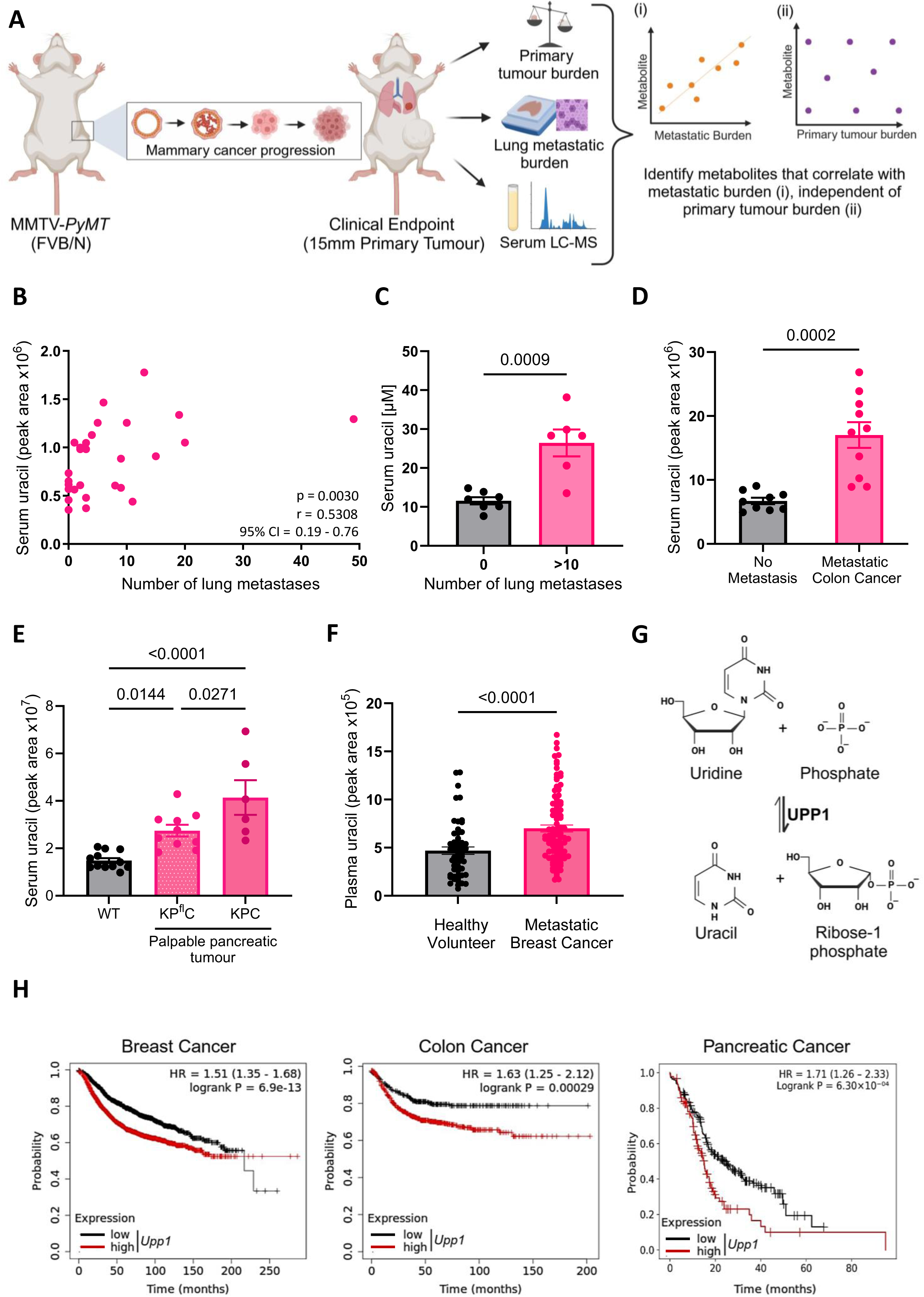
The UPP1 product uracil positively correlates with metastasis in mouse models of metastatic cancer. (A) Schematic representation of the experimental approach. MMTV-*PyMT* mice (n=29) were humanely culled when one primary tumour reached 15mm diameter. Primary tumour burden was expressed as a percentage of total body weight. Lungs were formalin-fixed and paraffin-embedded (FFPE), lungs were cut into a series of 4µm serial sections and metastatic burden assessed by manual assessment of hematoxylin and eosin (H&E) stain of sections 5, 10 and 15 using HALO imaging analysis software (Version v3.6.4134.137). Serum metabolites were profiled by LC-MS. Spearman rank correlations were calculated between metabolite peak area and metastatic burden, and metabolite peak area and primary tumour burden. Metabolites were identified that had significant correlations with metastatic burden (i) independent of any correlation with primary tumour burden (ii). Examples shown are pictorial representations of the trend of interest. (B) Serum uracil levels measured by LC-MS from the serum of MMTV-*PyMT* tumour bearing mice described in A, plotted against the number of lung metastases. Each dot represents an individual mouse (n=29). Spearman Correlation statistics are presented (p value, Spearman r and 95% confidence interval). (C) Absolute quantification of serum uracil levels in MMTV-*PyMT* mice with no identified lung metastases (n=7 mice) and MMTV-*PyMT* mice with >10 lung metastases (n=6 mice), using LC-MS and a standard curve of known concentrations of uracil, unpaired t-test, mean ± SEM. (D) Serum uracil levels were measured in the KPN mouse model of colon cancer, comparing serum from tail bleeds at 30 days post induction (no metastasis, n=9) to serum from mice at clinical endpoint (metastatic colon cancer, n=10), unpaired t-test, mean ± SEM. (E) Serum uracil levels were measured by LC-MS from KP^fl^C mice modelling poorly metastatic pancreatic cancer (n=10) and KPC mice modelling highly metastatic pancreatic cancer (n=6) at palpable pancreatic tumour, compared to age-matched wild-type (WT) controls (n=12), one-way ANOVA, mean ± SEM. (F) Uracil levels were determined by LC-MS from plasma of human metastatic breast cancer patients (n=99) and healthy volunteers (n=56), unpaired t-test, mean ± SEM. (G) Schematic representation of the enzymatic reaction catalysed by Uridine Phosphorylase 1 (UPP1). (H) Increased expression of *UPP1* correlates with decreased relapse free survival in human breast cancer and colon cancer, and decreased overall survival in pancreatic cancer (PDAC grade G1-G4, Stage S1-S4, T1-T4, N0-N1, M0-M1). High and low expression were defined by KM Plotter auto select best cutoff. The hazard ratio (HR) from the Cox model is presented, and the consequent log-rank p-value [28, 29]. For B-F, each dot represents an individual mouse, or human, as appropriate.

Uracil is one of the four nucleobases in RNA and, whilst pyrimidine nucleotides can be produced through *de novo* biosynthesis, uracil is a product of the pyrimidine salvage pathway, namely the UPP1, catalysed phosphorolysis of uridine into uracil and ribose-1 phosphate (Figure 1G). Indeed, mice which lack the gene for *Upp1 (Upp1^-/-^*-mice), display increased levels of uridine and decreased levels of uracil in tissues and serum (Figure S2A-B). Furthermore, in human cancer patients, increased expression of *UPP1,* in tumour core biopsies, consistently correlates with decreased survival in breast, colon and pancreatic cancer (Figure 1H) [28, 29]. Collectively these data suggests that UPP1-dependent generation of uracil is associated with metastasis, and poor prognosis, in several cancer types.

### Neutrophils are a source of UPP1 in metastatic cancer

Given the pivotal role of inflammation in facilitating progression of metastatic disease, we investigated the serum metabolome of mice injected with lipopolysaccharide (LPS), an established inflammatory trigger, to understand which circulating metabolites associated with metastatic cancer could be generated by activated immune cells. LPS administration led to a consistent increase in serum uracil, indicating that UPP1-activation occurs during mobilisation and/or activation of immune cells (Figure 2A; Figure S3A). To further define the immune cell source of UPP1, transgenic mice expressing the diphtheria toxin receptor (DTR) under control of the CD11b promoter (CD11b-DTR) were treated with diphtheria toxin (DT) to deplete CD11b expressing cells. As CD11b is a pan-myeloid marker, this enabled non-specific ablation of cells across the myeloid lineage. Depletion of myeloid cells resulted in a striking decrease in serum uracil (Figure S3B-C), suggesting that the myeloid compartment is responsible for a significant proportion of uracil detected in the circulation. To further understand the cell type responsible for inflammatory - and metastasis - associated uracil production, we quantified *Upp1* in flow cytometry-sorted immune cells. Consistent with our previous observations, LPS challenge potently increased expression of *Upp1* in neutrophils isolated from the bone marrow (BM), blood and spleen (Figure 2B; Figures S3D-E), whilst *Upp1* was not detectable in T-cells sorted from the same tissue. Notably, LPS-driven *Upp1* expression had a tendency to be greater in Ly6G-high (mature) than in Ly6G-intermediate (immature) neutrophils (Figure 2B; Figure S3D-E), suggesting that *Upp1* expression is upregulated during neutrophil maturation in addition to following neutrophil activation [30]. Given the pronounced neutrophilia associated with tumour progression in the MMTV-*PyMT* model of mammary cancer (Figure S3F), and the positive correlation between circulating neutrophil counts and metastatic burden in these mice (Figure S3G), we assessed *Upp1* expression in neutrophils from MMTV-*PyMT* mice bearing mammary tumours. Increased *Upp1* was detected in BM neutrophils from tumour-bearing MMTV-*PyMT* mice compared to non-tumour bearing FVB/N controls (Figure 2C). Consistently, analysis of publicly available data also indicated that tumour-resident neutrophils from MMTV-*PyMT* mice expressed more *Upp1* than neutrophils isolated from the mammary fat pad in non-tumour-bearing mice (Figure 2D-E). Furthermore, using Microenvironment Cell Populations-counter (MCP-counter) analysis to assess immune infiltrates in human tissue [31], the co-occurrence of increased *UPP1* expression and a neutrophil signature was also reflected in transcriptomic data from human colorectal tumours, whilst no significant correlations were associated between *UPP1* expression and T cell signatures (Figures S3H).

**Figure 2:**
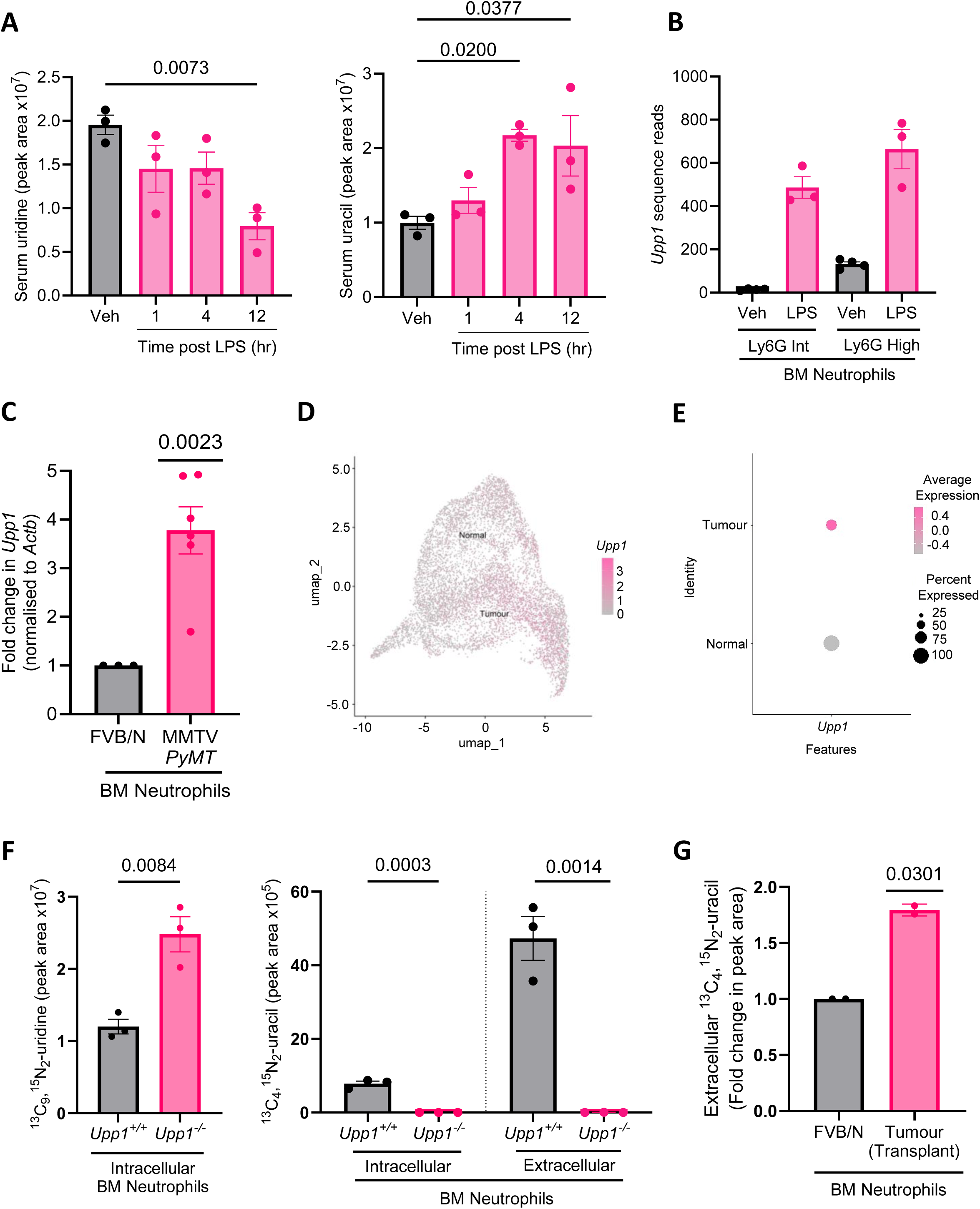
Neutrophils are a source of *Upp1*. (A) Wild-type (WT) mice on a C57BL/6 background were dosed intraperitoneal (IP) with 1mg/kg lipopolysaccharide (LPS), and serum uridine and uracil measured by LC-MS at defined timepoints post dosing, n=3 per experimental arm using male and female mice, one-way ANOVA, mean ± SEM. (B) *Upp1* transcript reads detected by RNA-Seq in CD11b^+^Ly6G^+^ neutrophils flow cytometry sorted from the bone marrow of C57BL/6 mice 24 hours post IP dosing of 1mg/kg LPS, one-way ANOVA, mean ± SEM, n=3-4 mice per experimental group from Mackey et al., 2021 [30]. Ly6G intermediate and high surface expression determined by flow cytometry was used to distinguish immature and mature neutrophils respectively. (C) *Upp1* was detected by qRT-PCR in neutrophils isolated from the bone marrow (BM) of female FVB/N (n=3) or tumour-bearing MMTV-*PyMT* mice (n=6). Data are presented as fold change compared to the age-matched WT mouse taken in parallel on the day of processing, one sample t-test assesses the MMTV-*PyMT* difference to 1, mean ± SEM. (D-E) scRNA-Seq data was analysed from GSE139125 [71] to assess *Upp1* expression in neutrophils isolated from the normal mammary fat pad versus neutrophils from *PyMT^+^* mammary tumour tissue. (F) Intracellular ^13^C_9_,^15^N_2_-uridine and intra-and extracellular ^13^C_4_,^15^N_2_-uracil were detected by LC-MS in the cells and conditioned media respectively from BM neutrophils isolated from female FVB/N *Upp1^+/+^* and *Upp1^-/-^* mice, that were incubated for 24 hours with ^13^C_9_,^15^N_2_-uridine, n=3 mice per experimental group, unpaired t-test, mean ± SEM. (G) Extracellular ^13^C_4_,^15^N_2_-uracil was detected by LC-MS from ^13^C_9_,^15^N_2_-uridine containing media conditioned by BM neutrophils isolated from female FVB/N mice and mice bearing 10–12mm tumours from mammary fat-pad transplantation of KP tumour fragments. Metabolites were harvested 24 hours post-plating, n=2 mice per experimental group. Data are presented as fold change compared to the WT mouse taken in parallel on day of processing. Mean ± SD, one sample t-test assesses the difference observed with BM neutrophils isolated from tumour-transplanted mice compared to 1. In all bar graphs each dot represents an individual mouse.

To interrogate the functional/enzymatic activity of UPP1 we isolated neutrophils from *Upp1^+/+^* and *Upp1^-/-^* mice, incubated them with ^13^C_9_,^15^N_2_-uridine, and used LC-MS to measure the ^13^C_4_,^15^N_2_-uracil produced from phosphorolysis of the isotopomer-labelled nucleoside. This indicated that UPP1 in neutrophils catalyses the phosphorolysis of uridine (Figure 2F, left-hand graph) and, moreover, that the uracil generated from this was exported into the medium (Figure 2F, right-hand graph). To assess the influence of a primary tumour on neutrophil UPP1 activity, mammary tumour fragments obtained from *K14*-Cre; *Trp53^fl/fl^* (KP) mice were orthotopically transplanted into the 4^th^ mammary fat pad of FVB/N recipient mice and allowed to grow to 10-12 mm in diameter. BM neutrophils were then isolated from tumour-bearing and non-tumour bearing FVB/N mice and their UPP1 activity determined by incubation with labelled uridine as previously described. This indicated that neutrophils from tumour-bearing mice had increased UPP1 activity, and consistent with the previous data, the increased uracil produced from UPP1-mediated cleavage of uridine was exported from the cell (Figure 2G). These data, therefore, indicate that both inflammatory triggers and the presence of primary tumours can not only drive the mobilisation of neutrophils, but can also increase *Upp1* expression within neutrophils, and suggests that neutrophils are a significant source of UPP1, and circulating uracil, in these pathologies.

### UPP1 influences neutrophil motility in the premetastatic lung through altered surface expression of CD11b

Several studies have described the importance of neutrophils in establishment of pre-metastatic niches, and how this contributes to metastasis of mammary cancer to the lung [7, 8]. More recent work has also described a slow-moving neutrophil population in the pre-metastatic lung of mice bearing mammary tumours [32]. Given the role of neutrophils in metastasis, and the upregulation of *Upp1* in neutrophils from mouse models of metastatic cancer, we sought to understand whether blocking UPP1 activity could influence neutrophil behaviour in the pre-metastatic lung. Firstly, we assessed the ability of 5-benzylacyclouridine (BAU), a competitive inhibitor of uridine phosphorylase, to block activity of UPP1 *in vivo*. Oral administration of 30mg/kg BAU, twice daily, suppressed uracil and elevated uridine levels, indicating effective inhibition of UPP1 *in vivo* (Figure S4A). Next, we orthotopically transplanted KP mammary tumour fragments into FVB/N recipients. Upon detection of a palpable mammary tumour, BAU was administered to mice orally, twice daily, and this treatment regimen was maintained until primary tumours grew to 10-12 mm in diameter – a timepoint in which neutrophils are mobilised by the primary tumour, but overt lung metastases are not yet detectable [32]. Because BAU did not influence primary tumour growth (Figure S4B), this provided the opportunity to assess the ability of UPP1 to influence the pre-metastatic microenvironment of the lung. Live cell imaging of precision-cut lung slices indicated that the migration speed of neutrophils was significantly decreased in tumour-bearing mice (Figure 3A-B), consistent with a previous report [32]. Strikingly however, inhibition of UPP1 with BAU completely opposed the ability of the primary tumour to decrease neutrophil motility in the pre-metastatic lung (Figure 3A-B), while BAU had no influence on neutrophil motility in the lungs of non-tumour bearing mice (Figure S4C). To understand how UPP1 might be influencing neutrophil migration we measured the expression of several cell surface molecules associated with neutrophil activation and adhesion (Table S2). This indicated that expression of the adhesion molecule CD11b (α_M_ integrin) was consistently increased on the surface of neutrophils isolated from lungs of mice bearing mammary tumours, which were either orthotopically transplanted (KP tumour) or autochthonous (MMTV-*PyMT*), by comparison with the CD11b surface expression detected in appropriate non-tumour bearing controls (Figure 3C-D). Importantly however, no increase in cell surface CD11b was detected in lung neutrophils from mammary tumour-bearing mice treated with BAU to inhibit UPP1 activity (Figure 3C-D; Figure S4D). To understand whether the UPP1-dependent increase in cell surface CD11b expression might be responsible for the decreased motility of neutrophils in the lungs of mammary tumour-bearing animals, we treated precision cut lung slices with M1/70, an antibody that blocks the interaction of CD11b with its ECM ligands. Treatment of precision cut lung slices, from mammary tumour-bearing mice, with M1/70 restored motility to lung neutrophils (Figure 3E). Collectively these data suggest that UPP1 can influence the surface expression of molecules such as CD11b, and the consequent cell surface availability of CD11b affects the migratory capacity of neutrophils in the pre-metastatic lung.

**Figure 3:**
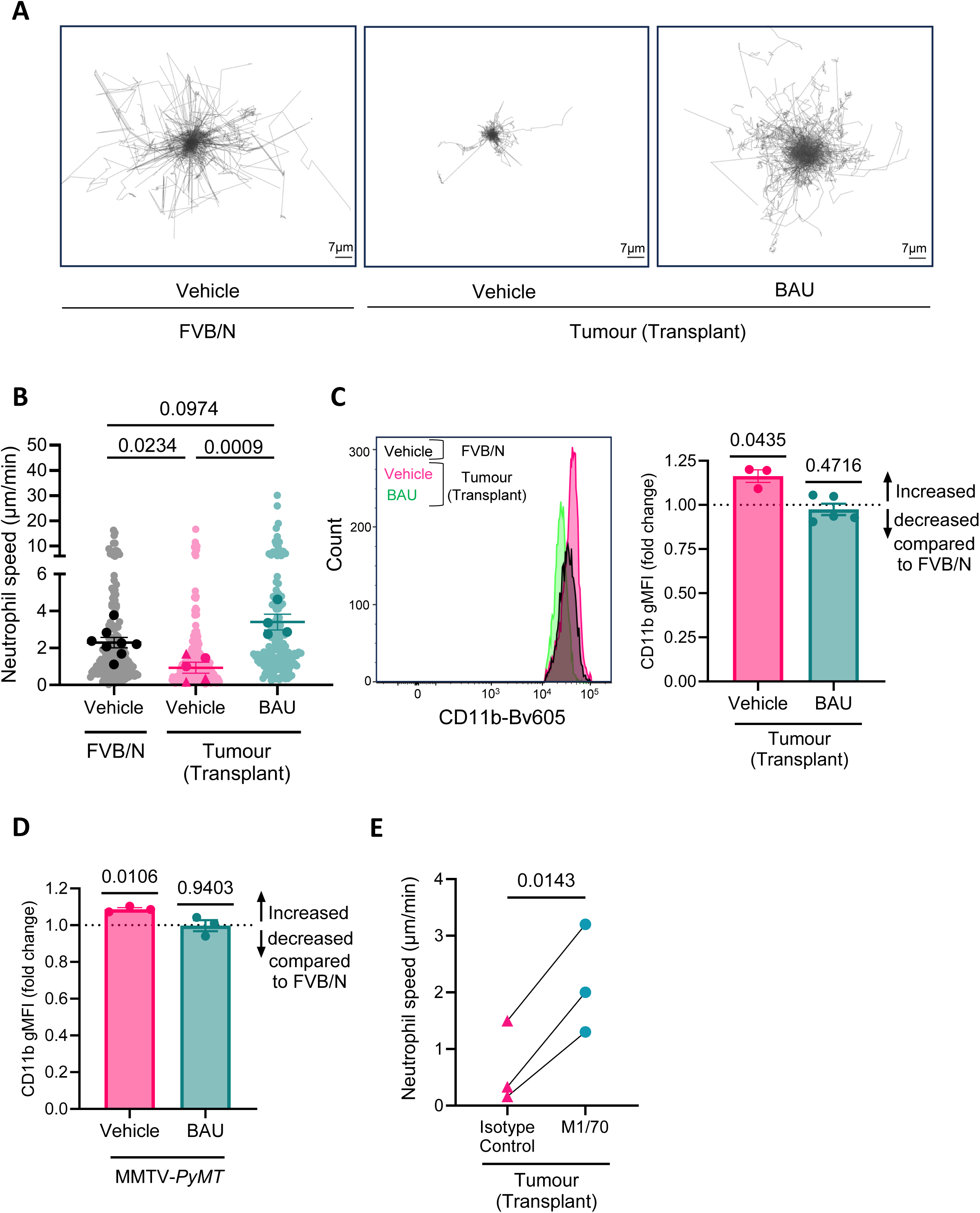
UPP1 influences neutrophil motility in the premetastatic lung through affecting surface expression of CD11b. (A) KP tumour fragments were transplanted into the 4^th^ mammary fat pad of female FVB/N mice. Upon detection of a palpable tumour mice were treated with vehicle (0.5% HPMC/0.1% Tween:DMSO, 90:10; p.o., b.i.d) or BAU (30 mg/kg p.o., b.i.d.). Once tumours reached 10 – 12mm in diameter, lungs were harvested, and the motility of neutrophils analysed by live cell imaging of precision cut lung slices (PCLS). Centred neutrophil tracks from moving neutrophils imaged by confocal timelapse microscopy of PCLS are presented, scale bars represent 7µm. (B) Motility of neutrophils from A quantified using Imaris software (Version 10.00). The motility of each individual moving neutrophil tracked is shown in lighter colours (up to 130 neutrophils per experiment) and the mean speed of neutrophil motility for each mouse is superimposed in darker colours, n=4-8 mice per experimental arm (n=8 vehicle treated FVB/N mice; n=5 vehicle treated KP tumour-transplanted mice, 3 of which had their lung slices treated with isotype control antibodies are presented in Fig 3E and are represented by triangles; n=4 BAU treated KP tumour transplanted mice). Statistics are performed on the mean speed per mouse data, one-way ANOVA, mean ± SEM. (C) Shifts in surface expression of CD11b on lung neutrophils were measured by quantifying the geometric median fluorescence intensity (gMFI) by flow cytometry of neutrophils (CD45^+^Ly6G^+^CD11b^+^) isolated from the lungs of FVB/N mice and FVB/N mice harbouring tumours from transplant of KP tumour fragments. Mice were treated as described in A, n=3 vehicle and n=5 BAU treated tumour-bearing mice. Because tumour-bearing mice were taken at a size-endpoint, an FVB/N mouse treated with vehicle for time-matched periods was taken in parallel on each day of processing to provide a baseline comparison. This resulted in n=6 FVB/N mice treated with vehicle. Data are presented as fold-change in CD11b gMFI in comparison to the FVB/N vehicle treated mouse taken in parallel (represented by the dotted line at 1). Statistics are a one sample t-test to assess whether each experimental group is different to 1, mean ± SEM. (D) CD11b measured as described in C, for MMTV-*PyMT* mice treated with vehicle or BAU from palpable tumour, n=3 mice in each experimental group, n=3 for FVB/N mice treated with vehicle to account of each day of processing. Statistics are a one sample t-test to assess whether each experimental group is different to 1, mean ± SEM. (E) Fold change in neutrophil speed measured by live cell imaging of PCLS from tumour-bearing vehicle treated mice described in A followed by ex vivo incubation with CD11b blocking antibody M1/70, or isotype control, n=3 per experimental group, paired t-test, mean ± SEM. In C – E, each shape represents an individual mouse, in E matched PCLS from n=3 mice are indicated by the adjoining line.

### UPP1-expressing neutrophils generate an immunosuppressed microenvironment in the lung

Neutrophils are capable of suppressing T cell proliferation, and this can generate immunosuppressed microenvironments that favour metastasis [33]. Given the accumulation of slow-moving neutrophils in the pre-metastatic lung (Figure 3 A-B) [32], we hypothesised that UPP1-driven alterations to the neutrophil phenotype might affect the landscape of other immune cell types within this metastatic target organ. Of all the cell populations assessed (Table S3), the most significant and consistent alterations following pharmacological inhibition or genetic deletion of *Upp1* were apparent in the numbers of T cells in the lung (Figure 4A-B; S5A-C). Specifically, treatment of MMTV-*PyMT* tumour-bearing mice with BAU led to an increase in the number of CD4^+^ and CD8^+^ T cells in the lungs (Figure 4A; Figure S5B). Consistently, increased numbers of CD8^+^ T cells were also observed in the lungs of MMTV-*PyMT* mice in which *Upp1* had been deleted (*Upp1*^-/-^ mice) (Figure 4B), corroborating the ability of UPP1 to influence T cell number in metastatic target organs. We also assessed T cell effector functions and, whilst no statistically significant alterations in the number of Granzyme B, IFN-γ or TNF-α positive CD8^+^ T cells were detected (Figure S6A-C), significantly increased numbers of IL-2^+^ CD8^+^ T cells were observed in the lungs of tumour-bearing MMTV-*PyMT* mice lacking *Upp1* (Figure 4C). IL-2 is a cytokine known to enhance T cell proliferation [34–37]. To understand whether alterations in local levels of UPP1 substrate and product, uridine and uracil respectively, could influence T cell proliferation, T cells were isolated from FVB/N mice, induced to proliferate by incubation with CD3/CD28 positive beads, and the ability of uridine or uracil to enhance or suppress proliferation was determined. This indicated that neither 30, 60 or 120µM uridine or uracil could alter the proliferation of T cells (Figure S6D). However, to study whether the physical presence of neutrophils might influence T cell proliferation in a way that was dependent on their UPP1-status, we measured T cell proliferation in the presence and absence of neutrophils from tumour-bearing MMTV-*PyMT* mice that were either *Upp1*^+/+^ or *Upp1*^-/-^ . This indicated that *Upp1*-expressing neutrophils from tumour-bearing mice strongly suppressed T cell proliferation, whilst neutrophils isolated from *Upp1* knockout tumour-bearing mice had a reduced ability to suppress T cell proliferation (Figure 4D-F). Taken together, these data suggest that the increased numbers of T cells in the lungs of mammary tumour-bearing *Upp1*-knockout mice may be accounted for by the reduced ability of *Upp1*^-/-^ lung neutrophils to suppress T-cell proliferation.

**Figure 4:**
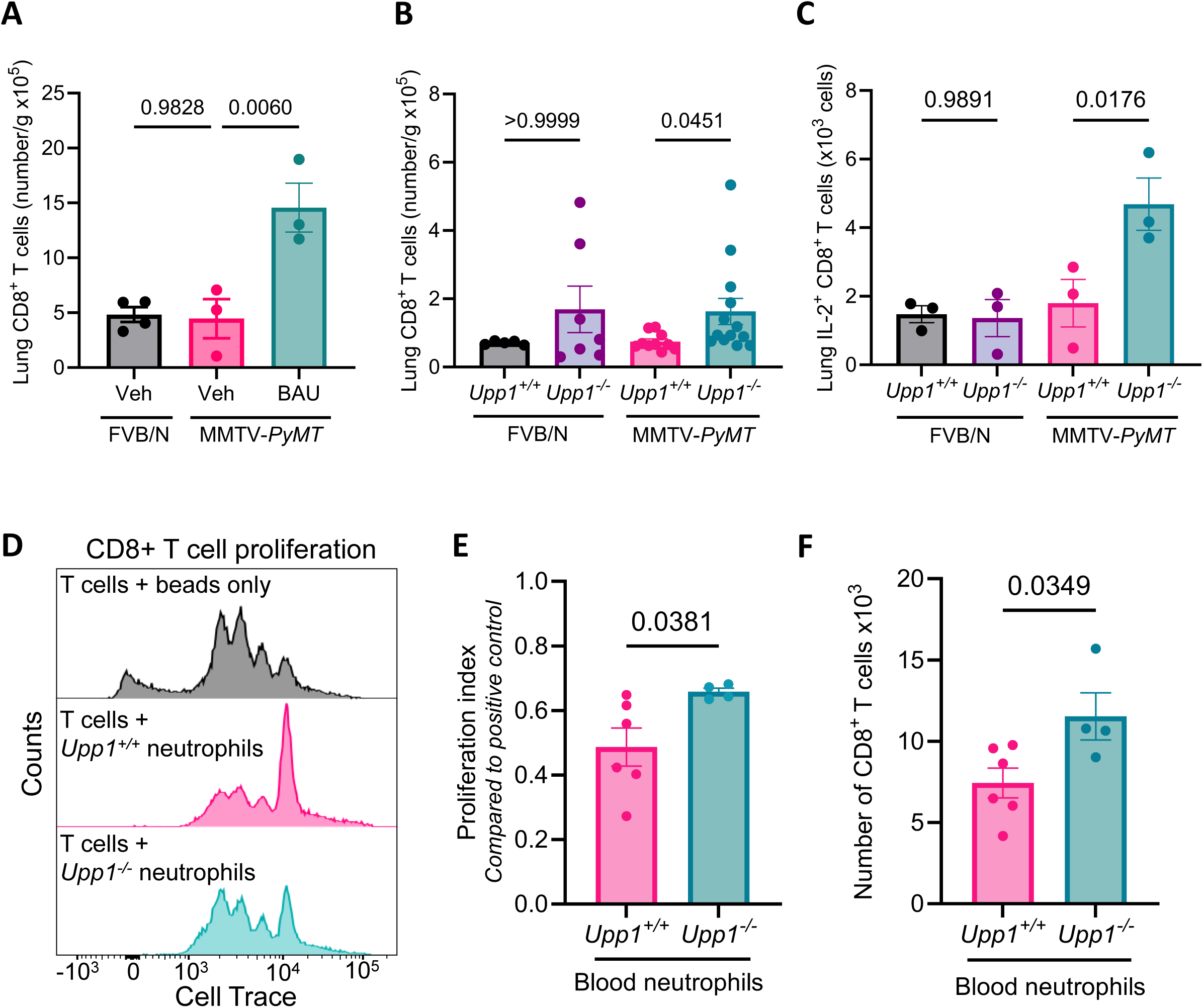
UPP1 influences the immune landscape of the pre-metastatic lung. (A) MMTV-*PyMT* mice were treated with vehicle (0.5% HPMC/0.1% Tween:DMSO, 90:10; p.o., b.i.d) or BAU (30 mg/kg p.o., b.i.d.) following detection of a palpable tumour, with FVB/N age-matched controls treated with vehicle for matched timepoints. Lungs of mice were harvested when one mammary tumour reached 10 – 12mm in diameter, and the number of CD8^+^ T cells assessed by flow cytometry, n=3 - 4 female mice per experimental group, with each mouse represented as an individual dot, one-way ANOVA, mean ± SEM. (B) Lungs of MMTV-*PyMT*;*Upp1^+/+^*and MMTV-*PyMT*;*Upp1^-/-^* mammary tumour-bearing mice were harvested when one tumour measured 10–15mm diameter and the number of CD8^+^ T cells was assessed by flow cytometry, n=5-13 female mice per experimental group with each mouse represented as an individual dot, one-way ANOVA, mean ± SEM. (C) Cells were prepared as B but with a 3 hour stimulation with PMA and ionomycin together with Brefeldin A, to enable assessment of intracellular interleukin 2 (IL-2) levels, n=3 female mice per experimental group, one-way ANOVA, mean ± SEM. (D) CD8^+^ T cells from the spleen and lymph nodes of FVB/N mice were incubated with equal numbers of neutrophils from the blood of tumour-bearing MMTV-*PyMT*;*Upp1^+/+^* and MMTV-*PyMT;Upp1^-/-^* mice and stimulated to proliferate with CD3/CD28 dynabeads for 48 hours. Representative histograms of cell divisions are shown, and quantified as (E) cell proliferation index and (F) total number of CD8^+^ T cells, as assessed by Cell Trace staining and flow cytometry, n=4–6 female mice per experimental group, with each mouse represented as an individual dot on graph, unpaired t-test, mean ± SEM.

### UPP1 activity can influence ECM deposition in the lung

In addition to altered immune landscapes, the deposition of ECM proteins is also considered to be a key feature of pre-metastatic niche priming. Increased levels of ECM components are reported to occur in several organs – including the lung and liver – prior to metastasis [38]. Indeed, increased fibronectin deposition in the lung was recently highlighted as a contributor to metastasis in mouse models of mammary cancer [39]. To understand whether UPP1 might influence the ECM-driven pre-metastatic niche priming, we used immunofluorescence to evaluate fibronectin deposition in the lungs of tumour-bearing mice that were either wild-type or knockout for *Upp1*. This indicated that lung fibronectin levels were significantly decreased in MMTV-*PyMT* tumour bearing mice in the absence of *Upp1* (Figure 5A-B).

**Figure 5:**
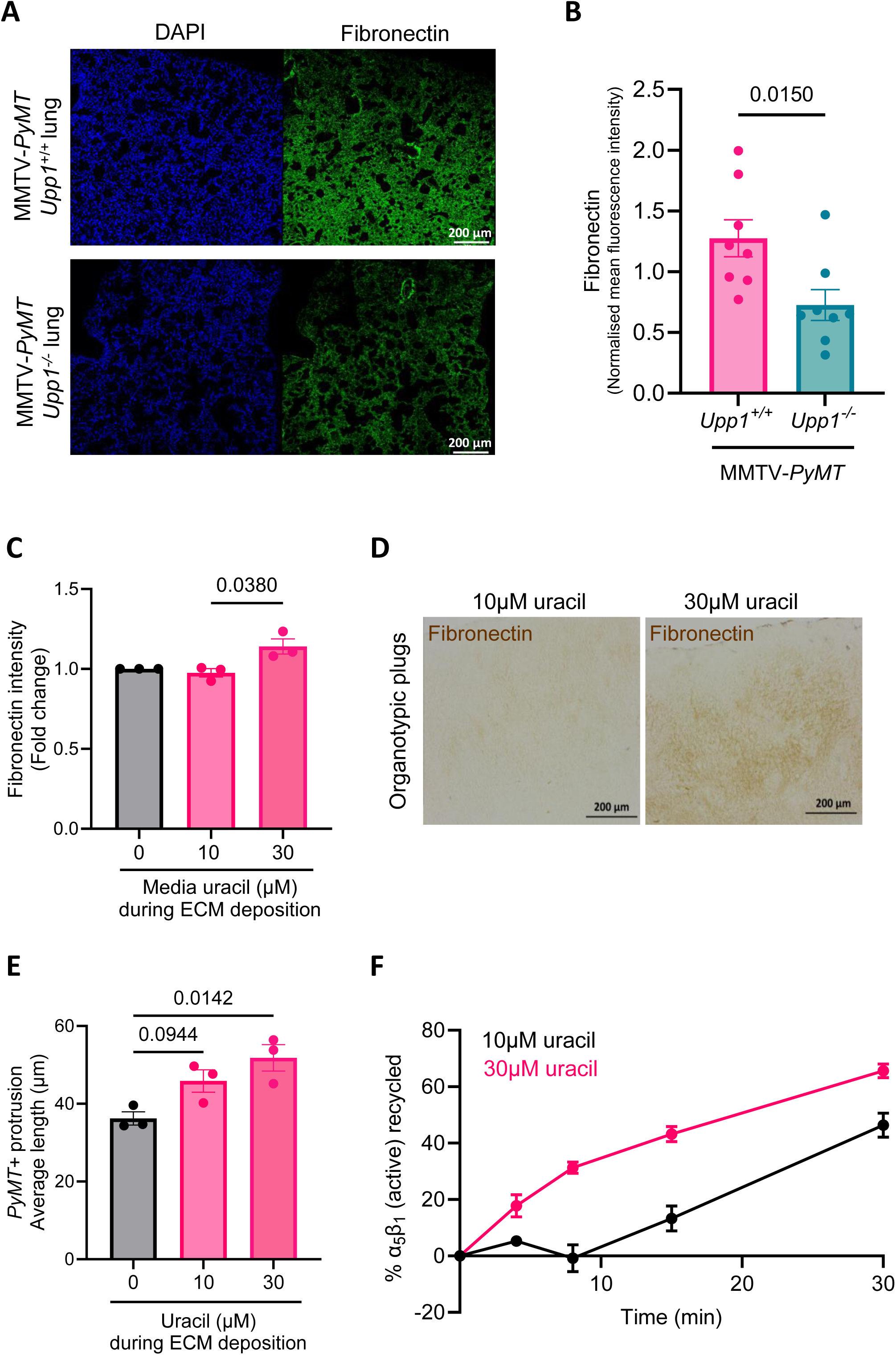
Extracellular uracil increases fibronectin deposition by stromal cells to generate a pro-migratory extracellular matrix. (A) Fibronectin in the lungs of clinical endpoint MMTV-*PyMT*;*Upp1^+/+^*and MMTV-*PyMT*;*Upp1^-/-^* mice was assessed by immunofluorescence. Representative images are shown from a total of n=8 female mice per experimental group that were assessed, scale bar is 200µm. (B) Quantification of lung fibronectin from mice described in A, with immunofluorescence quantified using FIJI (Version 2.9.0). Each dot represents an individual mouse at clinical endpoint, n=8 per experimental group, unpaired t-test, mean ± SEM. (C) Fibroblasts were plated to confluence and medium was supplemented with ascorbic acid and either vehicle, 10µM or 30µM uracil for 7 days. Fibroblasts were removed and the fibronectin content of deposited matrix was assessed by immunofluorescence staining and imaging on the Opera Phenix. The intensity of fibronectin was quantified using Columbus Software (Version 2.8.0), with the mean presented for n=3 biological repeat ECM preparations, normalised to the intensity of the matrix prepared in 0µM uracil, unpaired t-test, mean ± SEM. (D) Fibroblasts were added to rat tail collagen with medium containing 10µM or 30µM uracil. Collagen plugs were allowed to contract for 7 days, fixed in 4% PFA, embedded in paraffin and stained for fibronectin by immunohistochemistry. Representative images are shown for n=4 biological repeats, scale bar 200µm. (E) Pseudopod length of *PyMT*^+^ cancer cells migrating on cellular derived matrix was determined using ImageJ (Version 1.49). 10 cells were measured per field of view, for 6 positions per well. Each dot represents the mean pseudopod length per experiment for n=3 biological repeats, one-way ANOVA, mean ± SEM. (F) The recycling of the active confirmation of the fibronectin receptor, α_5_β_1_ integrin, was assessed in fibroblasts treated with 10µM or 30µM uracil for 24 hours. Data presented are the mean ± SEM from n=3 biological repeats.

Although fibronectin may be deposited by several cell types, fibroblasts are generally accepted to be a principal depositor of ECM proteins, and altered fibroblast phenotypes are a key niche priming event in metastatic target organs [40]. Given our previous data showing that UPP1-expressing neutrophils from tumour-bearing mice export uracil into the extracellular microenvironment (Figure 2G), and the consistent finding that increased uracil is detected in the circulation of mice and humans with metastatic cancer (Figure 1B-F), we hypothesised that increased extracellular uracil generated by UPP1-expressing cells may influence ECM deposition by fibroblasts. To test this, we treated fibroblasts with uracil and measured the fibronectin content of the ECM deposited by these cells. Addition of 30µM uracil (the concentration of uracil associated with high metastatic burden in MMTV-*PyMT* tumour bearing mice (Figure 1C)) enabled mouse fibroblasts to deposit an ECM with increased fibronectin content in both 2D and 3D microenvironments, whereas 10µM uracil (associated with little or no metastatic burden) was less effective in these regards (Figure 5C-D). Notably, these changes in fibronectin deposition were not accompanied by alterations in underlying fibrillar collagen deposition or organisation (Figure S7A-B). As fibronectin is a key contributor to the motility and invasiveness of many cell types including cancer cells, we allowed fibroblasts to deposit ECM in the presence and absence of extracellular uracil, de-cellularised the deposited ECM, and assessed the ability of this matrix to influence the migration of cancer cells (cells derived from *PyMT^+^* primary mammary tumours, named *PyMT^+^* cells). This indicated that the length of invasive protrusions extended by the cancer cells, a surrogate marker of their invasive capacity [41], was increased when they were plated onto ECM that had been deposited in the presence of increased extracellular uracil (Figure 5E).

The α_5_β_1_ integrin heterodimer is the cell’s main fibronectin receptor, and the behaviour of this integrin is key to deposition of fibronectin-containing ECM. The expression levels and ligand-binding affinity of α_5_β_1_ are important for its ability to drive fibronectin polymerisation. However, neither of these indices of α_5_β_1_ activation were influenced by addition of extracellular uracil to fibroblasts (Figure S7C-D). Integrin functions, and their ability to influence ECM deposition, are also controlled by the rates at which these ECM receptors are endocytosed and subsequently returned, or recycled, to the plasma membrane [42]. We, therefore, measured the rate at which internalised α_5_β_1_ was recycled from endosomes to the plasma membrane in the presence of increased extracellular uracil. This indicated that a concentration of uracil associated with high levels of metastasis (30µM) significantly increased the recycling of α_5_β_1_ (Figure 5F; Figure S7E). Moreover, uracil also increased recycling of other ‘cycling’ receptors, such as the transferrin receptor (Figure S7F) indicating that the ability of uracil to influence ECM deposition could be via general modulation of endosomal recycling and not specifically due to specific control of the α_5_β_1_ trafficking. And so, in addition to the influence of UPP1-status on neutrophil motility and immunosuppressive behaviour in the lung, we find that the increased extracellular uracil produced by these neutrophils can also have non-cell autonomous effects in the local microenvironment, collectively contributing to the establishment of microenvironments that would be likely to promote metastasis.

### UPP1 promotes lung metastasis in a mouse model of mammary cancer

Our data indicate that UPP1 controls neutrophil behaviour, T-cell number and ECM deposition in the lungs of tumour-bearing mice, suggesting that this this enzyme can make key contributions to the progression of metastatic disease [43]. The ability of UPP1 to influence progression of mammary cancer was therefore assessed by comparing the primary tumour development and incidence of lung metastasis in MMTV-*PyMT:Upp1^+/+^* mice to that in MMTV-*PyMT:Upp1^-/-^* mice. Whole-body genetic deletion of *Upp1* did not influence the time to clinical endpoint (one primary tumour reaching 15mm in diameter) (Figure 6A), nor did it influence the final primary tumour burden of MMTV-*PyMT* mice (Figure 6B; Figure S8A). However, the proportion of MMTV-*PyMT* mice that had lung metastases at clinical endpoint was significantly reduced in *Upp1*^-/-^ mice (Figure 6C; Figure S8B-D), indicating that the role of UPP1 in mammary cancer is specific to the metastatic process. Collectively, our data therefore suggests that UPP1 influences immune cell landscapes and ECM deposition in a metastatic target organ, and in the context of mammary cancer this manifests as an ability of UPP1 to promote lung metastasis (Figure 6D).

**Figure 6:**
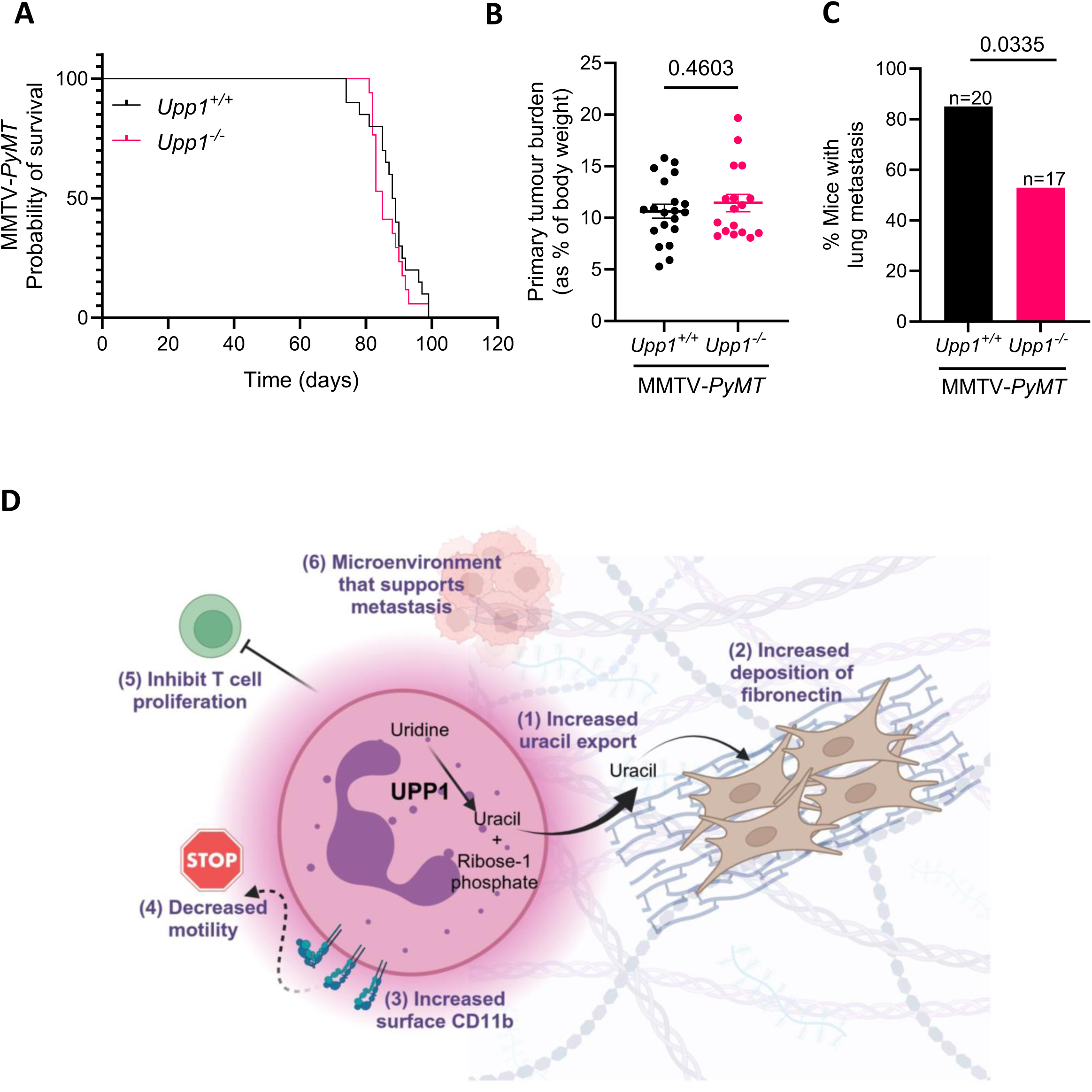
UPP1 specifically influences metastasis in a mouse model of mammary cancer. (A) Survival of MMTV-*PyMT*;*Upp1^+/+^*(n=20) and MMTV-*PyMT*;*Upp1^-/-^* (n=17) females, with mice culled at clinical endpoint, corresponding to one mammary tumour reached 15mm in diameter, log-rank (Mantel-Cox) test p>0.9999. (B) Primary tumour burden was measured by weight of total mammary tumours as a proportion of total mouse body weight, n=17–20 per experimental group as defined in A, unpaired t-test, mean ± SEM. (C) Lungs were formalin-fixed and paraffin-embedded (FFPE) and cut into a series of 4µm serial sections and metastatic burden assessed by manual assessment of hematoxylin and eosin (H&E) stain of sections 5, 10 and 15 using HALO imaging analysis software (Version v3.6.4134.137). The proportion of MMTV-*PyMT*;*Upp1^+/+^*(n=20) and MMTV-*PyMT*;*Upp1^-/-^* (n=17) female mice with detectable lung metastases was assessed, Chi-squared test. (D) Schematic representation of the current working model: Neutrophils increase expression of *Upp1* in response to the presence of a primary tumour. UPP1 high neutrophils secrete uracil into the extracellular microenvironment, and increased extracellular uracil promotes deposition of a fibronectin rich extracellular matrix (ECM). UPP1 high neutrophils have high cell surface CD11b, resulting in decreased neutrophil motility in the pre-metastatic lung. Congested neutrophils inhibit T cell proliferation. Collectively these mechanisms create niches that support metastasis. In the absence of UPP1 expression or activity, neutrophil motility is restored, fibronectin deposition is decreased, T cell numbers are increased, and the ability of mammary cancer cells to successfully colonise metastatic target organs such as the lung is reduced.

## Discussion

Altered metabolism is a hallmark of cancer recognised to play an important role in the progression of metastatic disease. UPP1, the enzyme responsible for the phosphorolysis of uridine into uracil and ribose-1 phosphate, is a key nucleotide metabolism enzyme that has been the subject of several papers in recent years, in part due to its consistent and strong relationship with decreased survival in several cancer types. The ability of cells to proliferate and migrate is dependent on the *de novo* generation of macromolecules, and targeting nucleic acid synthesis through nucleotide antimetabolite therapy represents the backbone of therapy for many cancers [44]. Indeed, it is this paradox that makes UPP1 an interesting enzyme to study given that it facilitates a catabolic reaction which disassembles nucleosides but has expression levels that are consistently upregulated in cancer – a disease canonically associated with increased formation of macromolecules rather than their disassembly. The upregulation of UPP1 in cancer cells has inspired studies focused on the tumour-intrinsic effects of UPP1. This includes studies in pancreatic cancer, where UPP1-dependent production of ribose-1 phosphate was found to fuel central carbon metabolism in nutrient-deprived conditions [45, 46]. Tumour-specific *UPP1* increases have also been associated with disease progression in lung adenocarcinoma [47]. In these instances, the authors showed that increased UPP1 expression drove primary tumour growth. We find that, although UPP1 is key to establishment of lung metastases in mammary cancer, this enzyme does not seem to contribute to growth of primary mammary tumours. This may indicate that the ability of UPP1 to influence primary tumour growth may be more apparent in cancers of the pancreas and lung compared to mammary cancer. However, given that our data suggest that UPP1’s influence on metastasis is most likely mediated via immune cells, its lack of influence on growth of MMTV-*PyMT*-driven mammary tumours is likely owing to the ‘immune cold’ properties of primary tumours in this model [48], by comparison with lung and PDAC models with established immune dependencies at the primary site [45, 47, 49].

Whilst LPS-driven systemic inflammatory responses are distinct from those evoked by the presence of a primary tumour, we demonstrate that a key hallmark of the metastasis-associated metabolome – elevated circulating uracil – is a common metabolic feature of both inflammation and metastasis. Importantly, the GEMMs employed in the present study all recapitulate metastasis through mechanisms driven by altered neutrophil biology [7–11]. We show here that neutrophils mobilised in response to the presence of a primary tumour generate and release uracil which makes a key contribution to the metastasis-associated circulating metabolome, and that this may influence ECM deposition by fibroblasts to help prime the lung metastatic niche. Importantly, activity of the enzyme responsible for producing uracil, UPP1, also influences neutrophil behaviour in the lung and the ability of these cells to affect T-cell proliferation – mechanisms that can generate immunosuppressed environments that are permissive to metastatic seeding and outgrowth. The ability of neutrophils to facilitate metastasis is well-established, and increased numbers of neutrophils in metastatic target organs results in immunosuppressed microenvironments that allow disseminated cancer cells to thrive [7–9]. Recent work has also highlighted differences in the movement of neutrophils in the pre-metastatic lung, with lung neutrophils having decreased motility in mice bearing primary mammary tumours [32]. The migration of leukocytes can be influenced by the cell surface expression of transmembrane receptors, including integrins. Indeed, integrins are known to influence cell adhesion and motility in a complex way. Very low levels of specific integrins at the cell surface are associated with decreased leukocyte migration due to a reduced ability of cells to engage the substratum [50]. However, high levels of cell surface integrin can also result in decreased motility due to excessive adhesion to the ECM [51, 52]. Thus, cells struggle to migrate when surface levels of integrin are either very high or very low, whilst intermediate levels are associated with efficient cell migration. We have found that the presence of a primary tumour in the mammary gland increases neutrophil CD11b surface expression, which stalls neutrophil migration in the lung. Moreover, this primary tumour-driven stalling of neutrophil motility is relieved by blocking engagement of CD11b with its ECM ligands. Expression of cell surface proteins can commonly be regulated post-transcriptionally to enable efficient cellular responses to the local microenvironment, and post-transcriptional regulation of integrins is commonly exerted via modulation of endo- and exocytosis [53]. As α_5_β_1_ integrin recycling in fibroblasts is promoted by addition of extracellular uracil (Figure 5F; Figure S7F), this sets a precedent for UPP1’s ability to regulate CD11b surface expression via modulation of endosomal trafficking. However, addition of extracellular uracil to neutrophils does not increase CD11b surface expression (Figure S4E). Thus, although the data presented here suggest that UPP1 can, via release of uracil from neutrophils, influence integrin function in fibroblasts, this is unlikely to be how it controls integrin trafficking within the neutrophils themselves. It is interesting, therefore, to speculate how UPP1 might exert post-transcriptional control over neutrophil integrin trafficking and function. Indeed, integrin function is known to be impacted by metabolic perturbations that compromise glycosylation [54], and CD11b is a glycoprotein, with N-linked mannose high glycan epitopes being a described feature of CD11b from human neutrophils [55]. Moreover, glycosylation of cell surface receptors decreases their rate of internalisation [56], and pathway analysis finds that endocytosis is enriched in UPP1 high cells [45] – perhaps as a means to compensate for the decreased internalisation of surface proteins. Furthermore, UPP1 expression positively correlates with mannose metabolism in situations defined by high immune infiltration [57]. UDP-N-acetylglucosamine (UDP-GlcNAc) is a key precursor for the synthesis of glycan oligosaccharides used for N-linked glycoprotein assembly and glycosylation of integrins, and UDP-GlcNAc synthesis requires UTP. It is thus interesting to speculate that the excessive consumption of uridine, due to increased UPP1 activity, may be a way in which UPP1 can impact the global glycosylation state of the cell. Thus, the ability of UPP1 to influence glycosylation states and how this, in turn, effects integrin trafficking and function will be an interesting hypothesis to investigate in the future.

In the absence of UPP1 expression/activity we show that the number of T cells in the lungs of mice bearing mammary tumours increases. Furthermore, the expanded T cell population in the lungs of mammary tumour-bearing *Upp1* knockout mice express IL-2, a cytokine that actively promotes T cell proliferation [58]. Indeed, whilst lung neutrophils from *Upp1^+/+^* tumour-bearing mice oppose T cell proliferation, a factor that contributes to immunosuppressed microenvironments, lung neutrophils from tumour-bearing *Upp1^-/-^* mice have a decreased ability to suppress the proliferation of cytotoxic CD8^+^ T cells, suggesting that UPP1 may influence the immunosuppressive capabilities of neutrophils. Suppression of anti-tumour leukocytes at metastatic target sites by myeloid cells may be mediated through several mechanisms, including via increased production of inducible nitric oxide synthase (iNOS) or reactive oxygen species (ROS) [12, 59]. The ability of myeloid cells to reduce T cell responses can also depend on other interactions with the local microenvironment. Indeed, fibronectin can polarise myeloid cells towards immunosuppressive states via direct engagement of the ITL3 receptor [60], and chronic stress was recently shown to increase lung metastasis through a mechanism involving fibronectin accumulation, neutrophilia, and reduced T cell numbers [39]. Furthermore, UPP1 inhibition has previously been shown to protect against generation of fibrotic microenvironments [61]. Thus, given our observations that fibronectin deposition in the lungs of tumour-bearing mice is reduced by *Upp1* knockout, and that the UPP1 product, uracil, promotes fibroblast-mediated fibronectin deposition, it will be interesting to determine the extent to which UPP1-dependent immune landscapes and ECM deposition combine to generate the immunosuppressed environment necessary to foster metastatic seeding in the lung.

Collectively our data show that neutrophils are a source of UPP1 in cancer, and that UPP1 can influence the immunosuppressive and motile behaviour of neutrophils and ECM deposition in the lung. Importantly, in the absence of UPP1 the ability of mammary tumours to metastasise is reduced, and we thus highlight a specific role for UPP1 in the metastatic process. The results of this study may have important therapeutic ramifications, including the likelihood that circulating uracil could be used as a biomarker to detect metastatic disease, whilst it also highlights the potential use of UPP1 inhibitors to decrease the risk of distant metastasis, and/or improve responses to cancer immunotherapy.

## Acknowledgments

We are grateful to all funders, described fully below, and the services and resources provided by the CRUK Scotland Institute. We express special acknowledgement to the Biological Services Unit, the Beatson Advanced Imaging Resource (BAIR), the Flow Cytometry Facility, the Histology Facility, Transgenic Technologies and the Metabolomics Facility of the CRUK Scotland Institute. We thank all members of the Norman lab for constructive feedback throughout the course of this project. Mouse embryonic fibroblasts were kindly provided by the lab of Professor Sara Zanivan, *PyMT^+^* cells from the lab of Professor Karen Blyth, advice regarding use of GEMMs of pancreatic cancer by Professor Jen Morton, and data analysis methods were developed by Dr Ryan Corbyn, all at the CRUK Scotland Institute. We also thank Drs. Billy Clark and Ann Hedley, former CRUK Scotland Institute, for their contributions to transcriptomic data production and analysis, and Professor Jos Jonkers (Netherlands Cancer Institute) for the KP mouse model. All illustrations were created using BioRender.com. We reserve special thanks to the patients who allowed their samples to be used for scientific gain, and the murine models used throughout the course of this work. This paper was critically reviewed by Catherine Winchester, and we thank her for her contribution and role in research integrity at the CRUK Scotland Institute. We dedicate this paper to anyone that has been affected by metastatic cancer, in the hope that discoveries like this may help others in the future.

## Data Availability

Upon publication, transcriptomic, metabolomic and flow cytometry data will be deposited in appropriate public repositories and accession numbers provided (ArrayExpress, MassIVE and FLOW repository respectively for CRUK Scotland Institute produced data. Metabolomic data produced at University of California Irvine and University of Chicago will be shared in Metabolomics Workbench). All other data will be made available by the corresponding authors at reasonable request.

## Funding

The work described in this manuscript was supported by Cancer Research UK core funding awarded to JCN (CRUK Core A18277) and the CRUK Scotland Institute (A31287), and a Pancreatic Cancer UK Research Innovation Grant awarded to CJC and JVV (2021RIF_22_Voorde). Additional funders that supported authors throughout the time they contributed to this manuscript are detailed below:

## CJC

Wellcome Trust (318046). <u>JCN:</u> CRUK (A18277 funding the work of CJC, DW, SF, ED, DN, JP & LEM, plus A24388 & A26892), MRC (MR/P01058X/1 funding the work of MM) and Breast Cancer Now (2011NovPR12 funding ED). <u>KB:</u> CRUK (A29799 funding SD and LEM). <u>LMC:</u> CRUK (A23983), Breast Cancer Now (2019DecPR1424). <u>SBC:</u> Breast Cancer Now (2019DecPhD1349) and CRUK (A25142) funding the work of RW. <u>EWR:</u> CRUK (A1920). <u>OJS:</u> CRUK (A25187, A22645, A23390, A21139, A25265, A25233, A25045, A24388, A26825 and A28223), Bloodwise (15041). <u>CWS:</u> Academy of Medical Sciences (SGL018\1021), CRUK (A26018), CSO (PCL/17/03 & SCAF/19/01), NHS Greater Glasgow & Clyde (GN21ON369), Tenovus (S21-10).

## Author contributions

<u>Conceptualisation:</u> CJC, JCN, DW, JVV, KB, OJS. <u>Investigation:</u> CJC, DW, JCN, JVV, SF, DSump, SD, ED, JBGM, AJM, FF, JP, CDC, EA, RW, LEM, GT, LND, JJAS, CAL, AM. <u>Formal analysis:</u> CJC, DW, JCN, JVV, DSump, MM, DN, KG, KLR, JP, RW, LND, JJAS, CAL, AM, CJH. <u>Resources:</u> DSump, DStrath, RJ, IRM, SD, KB, CN, LMC, AM, CJH, OJS. <u>Methodology:</u> DSump, AMc, FF, LMC. <u>Supervision</u> of contributing authors: CJC, JCN, CN, CJH, AM, JVV, DStrath, PD, CWS, LMC, SBC, KB, OJS. <u>Writi</u>n<u>g – original draft:</u> CJC. <u>Writi</u>n<u>g</u> <u>– review & editing:</u> All authors were given the opportunity to provide feedback. The following authors provided feedback that was incorporated by CJC: JCN, DW, EWR, DSump, RW, MM, ED, JVV, FF, LMC, SBC, KB, IRM, CWS.

## Declaration of interests

The authors declare no competing interests.

## Methods

### Mice

The *Upp1* knock-out mouse line was generated at the CRUK Scotland Institute by CRISPR gene editing technology (*Upp1* gene ENSMUSG00000020407/GRCm39:CM001004.3). Two CRISPR guides were identified which cut in the intronic sequence surrounding exon 4 of the *Upp1* gene and were demonstrated to have high cutting efficiency in vitro; GTAGGGCCCTTGACAATGTA, ATAAACGCAGGCCACTTGTA. Guide RNAs were generated in house using Highscribe™ T7 Quick High Yield RNA synthesis kit (New England Biolabs Inc.), Alt-R™ S.p. Cas9 Nuclease was purchased from (Integrated DNA Technologies UK., Inc). To generate the electroporation solution; the guide RNAs and spCas9 protein were combined in Optimem™ (Thermo Fisher Life Technologies., Ltd). Approximately 7 h after in vitro fertilization, the 1-cell stage embryos of 5-6 week old FVB/N mice (Charles River UK) were introduced into electroporation solution, and electroporated using a NEPA21 electroporator (Nepa Gene Co., Ltd). The following day, 2-cell embryos were transferred into the oviducts of pseudopregnant CD1 females (Charles River UK). Genotyping of subsequent pups was performed from ear samples by PCR using the primers F: GCAGAGTGGGCTAGCAACTA and R: CAGTTGCAAGCAGGGATCAT followed by Sanger sequencing.

Female mice were used in all experimental studies using mammary cancer model. To model mammary cancer MMTV-*PyMT* mice [22] were backcrossed >20 generations and maintained on an FVB/N background. *K14*-Cre; *Trp53*^fl/fl^ (KP) mice were gifted from Jos Jonkers [62]. These mice were backcrossed to an FVB/N background for 5 generations. For orthotopic transplantation models of mammary cancer, 1mm^3^ mammary tumour fragments obtained from KP mice were transplanted into the 4^th^ mammary fat pad of 10–12-week-old female recipient FVB/N mice (Charles River UK) using appropriate anaesthesia and analgesia as previously described [63]. Mice were monitored for tumour development 2 – 3 times per week. Once detected, tumour growth was monitored by calliper measurement 3 times per week. For all mammary models, endpoints (time or size) are described in appropriate figure legends. Clinical endpoint is defined as one mammary tumour reaching 15mm in diameter.

For colon cancer, KPN mice (*villin*Cre^ER^; *Kras*^G12D/+^; *Trp53*^fl/fl^; *R26*^N1icd/+^) on a C57BL/6 background described previously [10] were induced with a single injection of 2mg tamoxifen by intraperitoneal injection at an age of 6 to 12 weeks. Both sexes were used. Serum was isolated from tail bleeds at 30 days post induction to provide baseline levels of metabolites in the absence of metastasis. Clinical endpoint was defined as weight loss and/or hunching and/or cachexia and confirmed metastatic cancer upon dissection.

For pancreatic cancer, KPC (*Pdx1*-Cre; LSL-*Kras*^G12D/+^; LSL-*Trp53*^R172H/+^) mice were used to model metastatic pancreatic cancer, and KP^fl^C (*Pdx1*-Cre; LSL-*Kras*^G12D/+^; LSL-*Trp53*^fl/+^) were used to model pancreatic cancer that has a decreased propensity to metastasise [26]. These mice are maintained on a mixed strain background and both sexes were used. Mice were sampled when pancreatic tumours were detectable by palpation, a timepoint in which pro-metastatic structural changes are apparent in distant target organs but that precede the formation of overt metastases [27]. Average KP^fl^C age at death was 19.30 weeks, and KPC 17.30 weeks: Difference in means −1.705 ± 2.778, 95% confidence interval −7.663 to 4.253, P value 0.5492 (unpaired t-test). Only mice with confirmed pancreatic tumours upon dissection were included for downstream analysis.

For experiments utilising the CD11b-DTR mice, mice were produced as an F1 cross between B6.FVB-Tg(*ITGAM*-DTR/EGFP)34Lan/J mice and C57BL/6J mice (Jackson Laboratory). Male mice between 8-12 weeks of age were used experimentally. Orthotopic PDAC tumours were established from 5×10^4^ KPC-7940B PDAC cells injected into the pancreas in 50µL of 1:1 Matrigel-DMEM + 10%FBS (Corning) of F1 C57BL/6J CD11b-DTR mice and allowed to establish tumours for 14 days. Mice were then treated with diphtheria toxin (Enzo Life Science) at a concentration of 25 ng/g or vehicle by intraperitoneal injection. Mice were humanely euthanised 24 hours after treatment.

In experiments where the immune system was mobilised in the absence of cancer, mice were dosed intraperitoneal with 1mg/kg (C57BL/6) or 0.5mg/kg (FVB/N) lipopolysaccharide (LPS) for 4 – 24 hours, as defined in individual figure legends. Both sexes were used at an age of 8 – 12 weeks.

As per ARRIVE (Animal Research: Reporting of *In Vivo* Experiments) guidelines 2.0: In all instances the experimental groups of comparison are described in the figure legend and associated text. Sexes of mice used per model are described above, and experimental endpoints (time or size) are described in the appropriate figure legends. Experimental units are a single animal, and the n of mice is represented as dots within graphs and reported in the corresponding figure legend. Exclusion criteria in cancer models applied only to mice that did not have cancer upon dissection (for example, mass due to cyst instead of tumour). Sample sizes are described in appropriate figure legends and influenced by the power needed to understand the biological effect being measured. Mice were randomly allocated into treatment arms. Confounding factors such as order of treatment and measurements were minimised by delegation of these tasks to staff blinded to the experimental design. Samples were labelled during processing and analysis in a way that did not indicate the treatment group to the researcher. Unblinding was performed once the processing and analysis of samples was complete. Outcome measures, statistical methods, experimental animals, procedures and results are all described within the figures and appropriate legends.

Except for CD11b-DTR mice, all mice were housed at the CRUK Scotland Institute, with environmental proactive enrichment, and in a ventilated barrier facility (12-hour light/dark cycle). Procedures were performed in accordance with the UK Animal (Scientific Procedures) Act 1986, approved by the local animal welfare (AWERB) committee, and conducted under UK Home Office licences 70/8645, 60/4181, 70/8646, P72BA642F, and PP6345023. Genotypes of genetically engineered mouse models were confirmed by Transnetyx genotyping services. Experiments using CD11b-DTR mice were conducted at University of California, Irvine, approved by the Institutional Animal Care and Use Committee (IACUC) under protocol AUP-23-084.

### Plasma from metastatic breast cancer patients and healthy volunteers

Plasma samples from patients with metastatic breast cancer and female healthy volunteers were originally collected under a protocol approved by West of Scotland REC4 (reference 10/S0704/32). Analysis of these archival samples for the current research was approved by Office for Research Ethics Committees Northern Ireland HSC REC B (reference 17/NI/0228).

### Untargeted metabolomics of MMTV-PyMT serum

Polar metabolites were extracted as previously described in [64], and this methodology is thus shared under the Creative Commons Attribution 4.0 International (CC BY 4.0) license (https://creativecommons.org/licenses/by/4.0/). Briefly, polar metabolites were extracted from the serum by the addition of ice-cold extraction solution (1:50 dilution, 20% water, 50% methanol and 30% acetonitrile) followed by centrifugation at 16,000g for 10 min at 4°C. The metabolite extract (supernatant) was stored at −74 °C until LC–MS analysis. Metabolite extracts (5μL) were injected and separated using a ZIC-pHILIC guard-analytical column (SeQuant; 20 mm × 2.1 mm; SeQuant; 150 mm × 2.1 mm, 5 μm; Merck) on an Ultimate 3000 HPLC system (Thermo Fisher Scientific). Chromatographic separation was performed using a 30-min linear gradient starting with 20% ammonium carbonate (20mM, pH 9.2) and 80% acetonitrile, terminating at 20% acetonitrile at a constant flow rate of 100μL min^−1^. The column temperature was held at 45°C. An online Q Exactive Orbitrap mass spectrometer (Thermo Fisher Scientific) equipped with electrospray ionization was used for both metabolite profiling and metabolite identification (LC-MS). For profiling, the polarity switching mode was used with a resolution (RES) of 70,000 at 200 m/z to enable both positive and negative ions to be detected across the mass range (75 to 1,000 m/z, with automatic gain control (AGC) target of 1×10^6^ and maximal injection time (IT) of 250ms). Data-dependent fragmentation was performed to aid metabolite identification using a pooled sample comprised of a mixture of all extracts. The Q Exactive was operated in positive and negative polarity mode separately (35,000 RES, AGC target of 1×10^6^ and max IT of 100ms), and the 10 most abundant ions were chosen for fragmentation (minimum AGC target of 1×10^3^, AGC target of 1×10^5^, max IT of 100ms, 17,500 RES, stepped normalized collision energy of 25, 60 and 95, isolation width of 1m/z, dynamic exclusion of 15s and charge exclusion of >2) per survey scan. The pooled sample was also used as a quality control (QC) and was repeatedly measured throughout the acquisition of the randomised biological extracts. Untargeted metabolomics analysis was performed using Compound Discoverer software (Thermo Scientific v2.1). Retention times were aligned across all data files (maximum shift of 2min and mass tolerance of 5ppm). Unknown compound detection (minimum peak intensity of 1×10^6^) and grouping of compounds were performed (mass tolerance of 5ppm and retention time tolerance of 0.2min). Missing values were filled using the software’s ‘Fill Gap’ node (mass tolerance of 5ppm and signal/noise tolerance of 1.5). Data were corrected for batch effects using the QC replicate injection data and a QC-based area correction regression model (Linear, 50% QC coverage, 30% RSD). Compound identification was assigned by matching the mass and retention time of observed peaks to an in-house library generated using metabolite standards (mass tolerance of 5ppm and retention time tolerance of 0.5min) or by matching fragmentation spectra to mzCloud (www.mzcloud.org; precursor and fragment mass tolerance of 10ppm and match factor threshold of 50).

### Uracil quantification in MMTV-PyMT serum

Uracil was quantified in serum by the standard addition method. A uracil stock solution was used to spike serum (pool of experimental samples) during extraction, generating a six point calibration curve ranging from 5μM to 160μM (0.1μM to 3.2μM at 1:50 extraction ratio). The calibration curves and samples were analysed together with the LC-MS profiling method described above. Targeted quantification was performed using Tracefinder (4.1, Thermo Scientific), a linear calibration curve was constructed by plotting the peak area of uracil against the standard addition amount. This was then used to calculate the uracil concentration in the unspiked samples.

### Targeted metabolomics

Targeted metabolomics was performed using the same instrument setup as described above for the untargeted analysis with modifications to the chromatography. The linear gradient was shortened to 15 minutes and the flow was increased to 200μl min^−1^. The mass spectrometry detection was carried out with the same parameters as above over the shortened LC runtime. Polar metabolites were extracted from serum by the addition of ice-cold extraction solution (1:50 dilution, 20% water, 50% methanol and 30% acetonitrile) followed by centrifugation at 16,000g for 10 min at 4°C. In cases where metabolites were extracted from tissue, 25mg of tissue was used per mL of extraction solvent, samples were homogenised at 4°C (Precelly’s 24 homogeniser with homogenising tubes for hard tissue) followed by centrifugation at 16,000g for 10 min at 4°C. The metabolite extract (supernatant) was stored at −74 °C until LC–MS analysis.

Targeted metabolomics analysis was performed using Skyline [65], version 21.2.0.369. A transition list containing the compound names, preferred adduct/polarity and measured retention times (with 2-min retention time tolerance) was generated using reference standards on the same LC-MS method. Extracted ion chromatograms were generated for each compound and the corresponding chromatographic peaks were carefully inspected and integrated. Relative quantification between sample groups was performed.

### ^13^C_9_,^15^N_2_-uridine metabolic tracing

Neutrophils were isolated from the femurs of MMTV-*PyMT* tumour bearing mice using MojoSort Mouse Neutrophil Isolation Kit (Biolegend), with neutrophil retention via negative selection strategy. Neutrophils were plated at a density of 450,000 cells/well in a 6 well plate using Plasmax [66] as a physiologically relevant cell culture medium, including 30µM ^13^C_9_,^15^N_2_-uridine (Cambridge Isotope Laboratories Inc), or unlabelled equivalent (Sigma), for 24 hours at 37°C/5% CO_2_. Cells where then removed from the plates by manual pipetting and centrifuged at 400g 4°C for 5 minutes to isolate the cell and media fractions. Polar metabolites were extracted using ice-cold extraction solution (20% water, 50% methanol and 30% acetonitrile). For intracellular extractions, 100µL was used per 1.50×10^5^ cells. For extracellular extraction a 1:50 dilution of media was used into extraction solution. Solutions were vortexed, followed by centrifugation at 16,000g for 10 min at 4°C. The metabolite extracts (supernatant) were then stored at −74°C until LC–MS analysis. Targeted metabolomics was performed as described above. As neutrophils are semi-adherent, the exact number of suspension neutrophils harvested may have differed to the number plated. For this reason, intracellular values were normalised to the cell count recorded per individual sample prior to lysis in extraction solution (implemented as a relative fold change between samples to preserve the peak area).

### Serum metabolomics of CD11b-DTR mouse model

Quantification of metabolites in serum from CD11b-DTR animals was performed as previously described [67, 68]. Briefly, 7 libraries containing 149 metabolites were serially diluted in water from 5mM to 1µM concentration. These external standard pools were used to calibrate isotopically labelled internal standards and allow for quantification of metabolites where internal standards are not available (Table S7). Metabolites were then extracted from 5µL of external standard pools or biofluid samples using 45µL of 75:25:0.1 acetonitrile:methanol:formic acid extraction mix to which the isotopically labelled internal standards described in Table S7 were added. After addition of the extraction mix, samples were vortexed for 10min at 4°C and then centrifuged at 15,000rpm for 10min at 4°C to pellet insoluble material. The supernatant containing polar metabolites was then moved to sample vials for analysis by LC-MS as previously described [67, 68].

XCalibur 2.2 software (Thermo Fisher Scientific) was used for metabolite identification and quantification. Metabolite identification was performed using the external standard libraries as reference to confirm the m/z and retention time for each metabolite. For quantification, when isotopically labelled internal standards were available, the concentration of internal standard in the extraction mixture was first determined by referencing the peak area of the isotopically labelled metabolite to peak areas of the unlabelled metabolite in the external standard dilution series. Upon determining the internal standard concentrations in the extraction mix, the peak areas of unlabelled metabolites in the biofluid samples were compared to the peak area of the quantified isotopically labelled internal standard to determine the concentration of the metabolite in the biofluid sample. For quantification of metabolites when were isotopically labelled internal standards were not available, the peak area of the metabolite was normalized in each sample to the peak area of an isotopically labelled internal standard with similar retention time. The normalised peak area of the metabolite in the biofluid sample was then compared to the normalised peak areas of the metabolite in the external standard dilution series to interpolate the concentration of the metabolite in the biofluid sample.

### Histological assessment of fixed murine tissue

Formalin fixed paraffin embedded (FFPE) lungs were cut into a series of 4µm serial sections and metastatic burden assessed by manual assessment of H&E stain in sections 5, 10 and 15 (using HALO image analysis software v3.6.4134.137). For each mouse the average number of metastases identified across the 3 sections of the lung was recorded, in addition to the average metastatic burden across the 3 lung sections (defined as (area of lung covered by metastases / total lung area) x 1000).

### Live precision cut lung slice imaging

Live cell imaging of precision cut lung slices was performed as described previously [32]. Briefly, lungs were inflated with low melting point agarose (Sigma), placed into ice-cold PBS, and sliced into 300µm thick sections using a vibratome. Lung slices were stained directly in chamber wells by direct incubation with Hoechst (ThermoFisher), Ly6G-AF647(1:200, Biolegend), CD31-AF488 (1:200, Biologend) (diluted in phenol-red free DMEM with 5% FBS) and imaged on a Zeiss LSM880 confocal microscope at 37°C/5%CO_2_ for 20 minutes with z-stacks of ∼20µm. Acquisition was performed with a 32 channel Gallium arsenide phosphide (GaAsP) spectral detector plan Apo objective lenses (20X/0,8 NA -dry, Carl Zeiss). Samples were excited simultaneously with 397 405, 488, 561 and 633 laser lines, with signal collected onto a linear array of the 32 GaAsp detectors in lambda mode with a resolution of 8.9 nm over the visible spectrum. Spectral images were then unmixed with Zen software (Carl Zeiss) using reference spectra acquired from unstained tissues (tissue autofluorescence) or slides containing single fluorophores. Timelapse image analysis was performed using Imaris (Bitplane). Neutrophils were segmented and tracked using the surface tool on Ly6G fluorescently labelled neutrophils, with surface detection and tracks assessed manually to ensure accuracy. In instances where the influence of the M1/70 blocking antibody on cell motility was assessed, 10µg M1/70 or isotype control (ThermoFisher) was added into the medium of the chamber-well.

### Flow cytometry

Lungs were mechanically dissected using dissecting scissors and shaking in RPMI medium (ThermoFisher) supplemented with 5% FBS, at 37°C for 20 minutes. Lung suspensions were filtered through a 70µm cell strainer, rinsed with RPMI medium, centrifuged at 400g 4°C for 5 minutes, and resuspended in RPMI medium supplemented with 5% FBS and 2mM EDTA (ThermoFisher). Cells were transferred to a 96 well v-bottomed plate for staining, using 50% of the final resuspended volume per stain. Briefly, cells were pelleted, washed in PBS, and resuspended in live/dead stain at room temperature protected from the light for 20 minutes. Cells were then washed in PBS, pelleted, resuspended in Fc Block and incubated at room temperature for 15 minutes. Cells were then stained with the relevant antibody cocktail (room temperature for 15 minutes in the dark), followed by sequential washing and fixation in PFA. Neutrophils were defined as Dead/Dump negative>CD45^+^>Ly6G^+^CD11b^+^, using the antibody panel described for neutrophil characterisation (Table S2). Total cell characterisation was performed using the antibody panel described in Table S3. T cells were further characterised by a lymphoid antibody panel (Table S4). Fluorescence minus one (FMO) controls were prepared to set gates. Count beads were added to samples to enable accurate determination of cell numbers. Data were acquired on the BD Fortessa (BD Biosciences) and analysed using FlowJo (BD Biosciences v10.9.0).

### Intracellular T cell staining

Lungs were mechanically dissociated with dissecting scissors and transferred into DMEM medium (ThermoFisher) supplemented with 1 mg/mL collagenase D (Roche) and 25 µg/mL DNase 1 (ThermoFisher). Enzymatic dissociation was carried out using the gentleMACS Octo Dissociator, run: 37C_m_LDK_01 (Miltenyi Biotec), as described [69]. Consequent lung suspensions were then filtered through a 70µm cell strainer, and enzyme activity was quenched by addition of FCS followed complete DMEM medium (supplemented with 10% FCS, 2 mM L-glutamine (ThermoFisher) and 10000 U/mL penicillin/streptomycin (ThermoFisher)). Cell pellets were lysed of red blood cells using commercially available 1x Red Blood Cell Lysis buffer (ThermoFisher), and following resuspension in PBS containing 0.5% BSA, cell number was acquired. After 3-hour stimulation with PMA and ionomycin, together with Brefeldin A (BioLegend), single-cell suspensions were incubated for Fc block (BioLegend) in 0.5% BSA/PBS buffer for 20 min on ice. Cell surface antibodies were added for 30 min at 4°C in the dark. Cell surface molecules were stained, and dead cells were identified with Zombie Green or Zombie NIR viability dye (BioLegend). Intracellular staining was performed after fixation and permeabilization with staining kit (00-5523-00, eBioscience). All antibodies used are described in Table S5. Gating strategy was Dead/Dump(CD19, CD11b and CD11c) negative>CD3^+^>CD8^+^>iL2^+^, IFNγ^+^, Granzyme B^+^ or TNFα^+^ as appropriate. Fluorescence minus one (FMO) controls were prepared to set gates. Count beads were added to samples to enable accurate determination of cell numbers. Data were acquired on the BD Fortessa (BD Biosciences) and analysed using FlowJo (BD Biosciences v10.9.0).

### T cell proliferation

Neutrophils were isolated from the blood of MMTV-*PyMT* tumour bearing mice using MojoSort Mouse Neutrophil Isolation Kit (Biolegend), with neutrophil retention via negative selection strategy. T cells were isolated from the spleen and lymph nodes of FVB/N mice using the MojoSort Mouse CD8 T cell isolation kit (BioLegend) as per the manufacturer’s protocol for negative selection strategy. T cells were stained with CellTrace Yellow Proliferation kit (Thermo Fisher Scientific). A total of 200,000 T cells were placed into a well of a flat-bottomed 96-well plate, and co-cultured with 200,000 neutrophils in 250μL Iscove’s modified Dulbecco’s medium (ThermoFisher), 10% FBS, 50mmol/L 2-mercaptoethanol (Sigma) with 2mmol/L L-glutamine (ThermoFisher) and 100U/mL penicillin-streptomycin (ThermoFisher). T cell proliferation was stimulated by addition of CD3/CD28 dynabeads at a ratio of 1 cell per bead (Gibco). Cells were harvested following 48 hours of incubation at 37°C/5%CO_2_ and assessed by flow cytometry, using the antibody panel described in Table S5.

### RNA-Sequencing

RNA-sequencing on flow cytometry sorted immune cell populations from vehicle and LPS treated mice was performed by Mackey *et al.* [30]. RNA-sequencing on mouse embryonic fibroblasts (MEFs) treated with uracil was performed as follows: MEFs were plated at 1×10^6^ cells per 10cm^2^ dish in full growth media (DMEM (ThermoFisher) supplemented with 10% FBS, 2mM L-glutamine (ThermoFisher) and 100U/mL penicillin/streptomycin (ThermoFisher)) supplemented with 10 or 30µM uracil (Sigma) and incubated at 37°C/5% CO_2_ for 24 hours. Cells were then washed with ice cold PBS, scraped into Trizol (ThermoFisher), and RNA prepped as per manufacturers instructions. RNA quality was assessed using Agilent 2200 Tapestation using RNA screentape. Libraries for cluster generation and DNA sequencing were prepared using Illumina TruSeq Stranded Total RNA Lib Prep, Gold kit. Quality and quantity of the DNA libraries was assessed on an Agilent 2200 Tapestation (D1000 screentape) and Qubit (Thermo Fisher Scientific) respectively. The libraries were sequenced on the Illumina Next Seq 500 using the High Output v2.5, 150 cycles kit (2 x 75cycles, paired end reads, dual index). N=6 biological replicates were prepared for each condition, however 1 sample from the vehicle and 1 sample from the 30µM uracil arm were removed at the data analysis stage due to human RNA contamination. Normalised read counts are presented.

### qRT-PCR

Neutrophils were isolated from the femurs of mice using MojoSort Mouse Neutrophil Isolation Kit (Biolegend), with neutrophil retention via negative selection strategy. Cells were collected into Trizol (Life Tech), and RNA isolated as per the manufacturer’s instructions. RNA was quantified and 260/280 ratio checked using Nanodrop (ThermoFisher). cDNA was synthesised using 1µg RNA as template using Quantitect Reverse Transcription Kit (Qiagen) as per the manufacturer’s instructions. qPCR was performed using x1 SyBr green (Qiagen), 1:10 dilution of cDNA template, 0.5µM forward primer (ThermoFisher), 0.5µM reverse primer (ThermoFisher), and reactions conducted using a 3 step protocol, corresponding to 95°C for 3 mins, x40 cycles of 95°C for 20 sec, 60°C for 20sec, 72°C for 20sec, with a final extension of 72°C for 5 mins, and melt-curve from 65°C to 95°C in 0.5°C increments. Data were collected and analysed in CFX Manager Software (BioRad CFX Maestro 1.0 Version 4.0.2325.0418). Expression of genes of interested was normalised to *Actb*. Primer sequences are described in Table S6.

### Cell derived matrices

Cell culture dishes were coated in 0.2% gelatin (Sigma), cross linked with 1% glutaraldehyde (Sigma), and quenched with 1M glycine (Sigma). Dishes were then washed with PBS and incubated with cell growth medium (DMEM (ThermoFisher) supplemented with 10% FBS, 2mM L-glutamine (ThermoFisher) and 100U/mL penicillin/streptomycin (ThermoFisher)) before addition of mouse embryonic fibroblasts at a concentration of 2×10^5^ cells per well in a 6 well plate. The following day, or when cells were deemed 100% confluent, medium was changed to full growth medium supplemented with 50µg/mL ascorbic acid (Sigma), plus 0, 10 or 30µM Uracil (Sigma) as per described experimental conditions, and medium was replaced and refreshed every other day for 7 days. Cells were incubated at 37°C/5%CO_2_ during the 7-day ECM deposition period. Cells were then washed with D-PBS with calcium and magnesium (named D-PBS from here on in, Sigma) and cells then removed by incubation with pre-warmed extraction buffer (20mM NH_4_OH, 0.5% Triton X-100 in D-PBS (Sigma)). Residual DNA was digested with 10µg/mL DNase I (Roche) and denuded matrices were then washed and stored in D-PBS with 100U/mL penicillin/streptomycin (ThermoFisher) at 4°C until use.

### Timelapse microscopy

Primary tumour cells derived from MMTV-*PyMT* mammary tumours (*PyMT^+^* cells) were plated at a density of 80,000 cells per well, in a 6 well plate, and allowed to adhere for ∼5 hours at 37°C/5%CO_2_. Cells were then imaged by timelapse microscopy, at 37°C/5%CO_2_, using an inverted phase-contract microscope (Axiovert S100, Carl Zeiss, 10x objective) taking images in 6 positions per well every 10 minutes for 16 hours. Images were sequentially processed to generate timelapse movies, and assessment of pseudopod length was performed using ImageJ software (Version 1.49).

### Immunofluorescence

For immunofluorescence of fibronectin in FFPE lung, antigen retrieval was performed by boiling in citrate buffer (10mM Citric Acid, pH 6, 0.05% Tween) for 25 minutes, followed by blocking in 1% BSA for 1 hour at room temperature and then incubation with rabbit polyclonal anti-Fibronectin (Abcam ab2413, 1:100) diluted in antibody diluent (Life technologies) overnight at 4°C. Fibronectin was detected using a Tyramide Signal Amplification (TSA-FITC) system (Perkin Elmer). Images were acquired on a Zeiss 710 using a 20x objective, with fibronectin intensity in the lung parenchyma quantified using FIJI software (Version 2.9.0).

For immunofluorescence of cell derived matrices (CDMs), CDMs were derived as previously described, in glass bottomed dishes (MatTek). Immediately after production, CDMs were fixed in 4% PFA, blocked in 1% BSA, and incubated in primary antibody for 2 hours at room temperature (Mouse anti fibronectin: BD Pharmingen 610078, diluted 1:100 in blocking buffer). Following washing, CDMs were then incubated with Alexa-555 conjugated secondary antibody (Themo Fisher 1:400). CDMs were imaged using the Opera Phenix High Content Screening System (Harmony High-content Imaging and analysis software version 4.9, PerkinElmer). Fibronectin fibres were identified and quantified using Columbus Image Data Storage and Analysis System (PerkinElmer version 2.8.0).

### Organotypic collagen plugs

Organotypic plugs were generated from rat tail-derived collagen as previously described [70]. Briefly, 2×10^6^ mouse embryonic fibroblasts were added to rat tail collagen in minimal essential medium (ThermoFisher) that had the pH adjusted to slightly acidic conditions, with either 10 or 30µM uracil (Sigma) as defined per experimental group. Collagen plugs were set at 37°C/5%CO_2_ for 10 minutes, and then fibroblast growth medium was added (DMEM medium supplemented with 10% FCS, 2 mM L-glutamine (ThermoFisher) and 100U/mL penicillin/streptomycin (ThermoFisher)), supplemented with 10 or 30µM uracil (Sigma). Plugs were kept at 37°C/5%CO_2_ and medium was refreshed every other day for 7 days. Plugs were cut in half, fixed in 4% PFA, embedded in paraffin in cross section, and stained using immunohistochemistry techniques. Fibronectin (ab2413, Abcam) was stained on a Leica Bond Rx, with antigen retrieval using ER2 solution (Leica) for 20 minutes at 100°C, and sections were rinsed with Leica wash buffer before peroxidase block was performed using an Intense R kit (Leica). To complete IHC staining sections were rinsed in tap water, dehydrated through graded ethanol’s, placed in xylene and then mounted onto coverslips in xylene using DPX mountant (CellPath). Fibrillar collagen in fresh organotypic plugs was visualised using a Trimscope multiphoton microscope (Lavision) equipped with a Ti:Sapphire laser at wavelength of 850 nm. For each field of view, Z-projections were acquired, and threshold was applied to the signal. The area of second harmonic generation (SHG) coverage per field of view was determined using ImageJ (Version 1.49). To assess organisation of fibrillar collagen in the plugs, we applied grey level co-occurrence matrix (GLCM), and focused on the parameter correlation. This texture analysis method determines the probability of pixels at increasing distances having similar intensities. Thus, if an image consists mainly of anisotropic fibres, it is possible to travel in a straight line away from a given point without much alteration to intensity, and this would give raise to longer decay distances for a given intensity correlation, in comparison to images containing isotropic fibres. For fields of view with unambiguous two-phase decay curves, the mean decay distance was calculated using method as described in [27].

### Receptor trafficking

Cells were plated in full growth medium supplemented with either 10 or 30µM uracil (Sigma) for 24 hours at 37°C/5%CO_2_. Medium was then removed, cells transferred to ice, washed twice in ice cold PBS, and surface-labelled at 4 C with 0.2 mg/mL NHS-SS-biotin (Pierce) in PBS for 30 min. Cells were transferred to full growth medium for 30 min at 37°C/5%CO_2_ to allow internalisation of the tracer. Cells were then returned to ice, washed twice with ice-cold PBS and biotin was removed from proteins remaining at the cell surface by reduction with MesNa (Fluka). The internalised fraction was then chased from the cells by returning them to 37°C/5%CO_2_ in full growth medium. At the indicated times, cells were returned to ice and biotin removed from recycled proteins by a second reduction with MesNa (Fluka). Biotinylated α_5_β_1_ was then determined by capture-ELISA using Maxisorp (Nunc) plates coated with antibodies specific to CD29 (Clone 9EG7, BD Pharmingen 550531) for active α_5_β_1,_ CD49e (Clone 5H10-27, BD Pharmingen 553319) for total α_5_β_1_, and transferrin receptor (BD Pharmingen 553264).

### Computational approaches

Survival curves presented in Figure 1 were generated using KM Plotter [28]. Normalised sequence reads for *Upp1* presented in Figure 2B, and Supplementary Figure 3D-E are from the flow cytometry sorted cells and transcriptomics described in [30]. Supplementary Figures 3H uses a Microenvironment Cell Populations-counter (MCP method) [31] on GSE33113. For the analysis of publicly available scRNA-Seq data (Figure 2D-E), data from (GSE139125) from mouse normal and PyMT mammary cancer tissue was utilised [71]. Briefly, data were processed using Seurat (version 5.1.0) on R (version 4.4.0) and integrated by reciprocal principal component analysis (RPCA) using the IntegrateData function then scaled and normalized. Dimension reduction was performed using principal component analysis (PCA) followed by clustering using the FindNeighbors and FindClusters functions. Marker genes for individual clusters were determined using the FindAllMarkers function. Neutrophils were isolated by the authors by isolating clusters positive for marker genes *Ly6g* and *Cxcr2*.

### Statistics

All statistical analyses were performed in GraphPad Prism (Version 10.1.2). Correlations were plotted using Spearman’s rank correlation coefficient, which measures the strength and direction of the monotonic relationship between two variables. Unpaired t-tests were used when considering two independent populations of equal variance. Ordinary one-way ANOVA with Šídák’s multiple comparisons test was used when making comparisons across multiple independent populations of equal variance. When experimental units were harvested on different days (such as mice bearing tumours at specific size endpoints), data are presented as a fold-change compared to an appropriate control sampled in parallel on the same day. In these instances, one sample t-tests have been used to assess whether the experimental group is different to 1. Paired t-tests were used on paired biological samples subjected to different treatments. All n are represented as dots on graphs, and reported numerically in the corresponding figure legends.

---

**Supplementary Figure 1:**
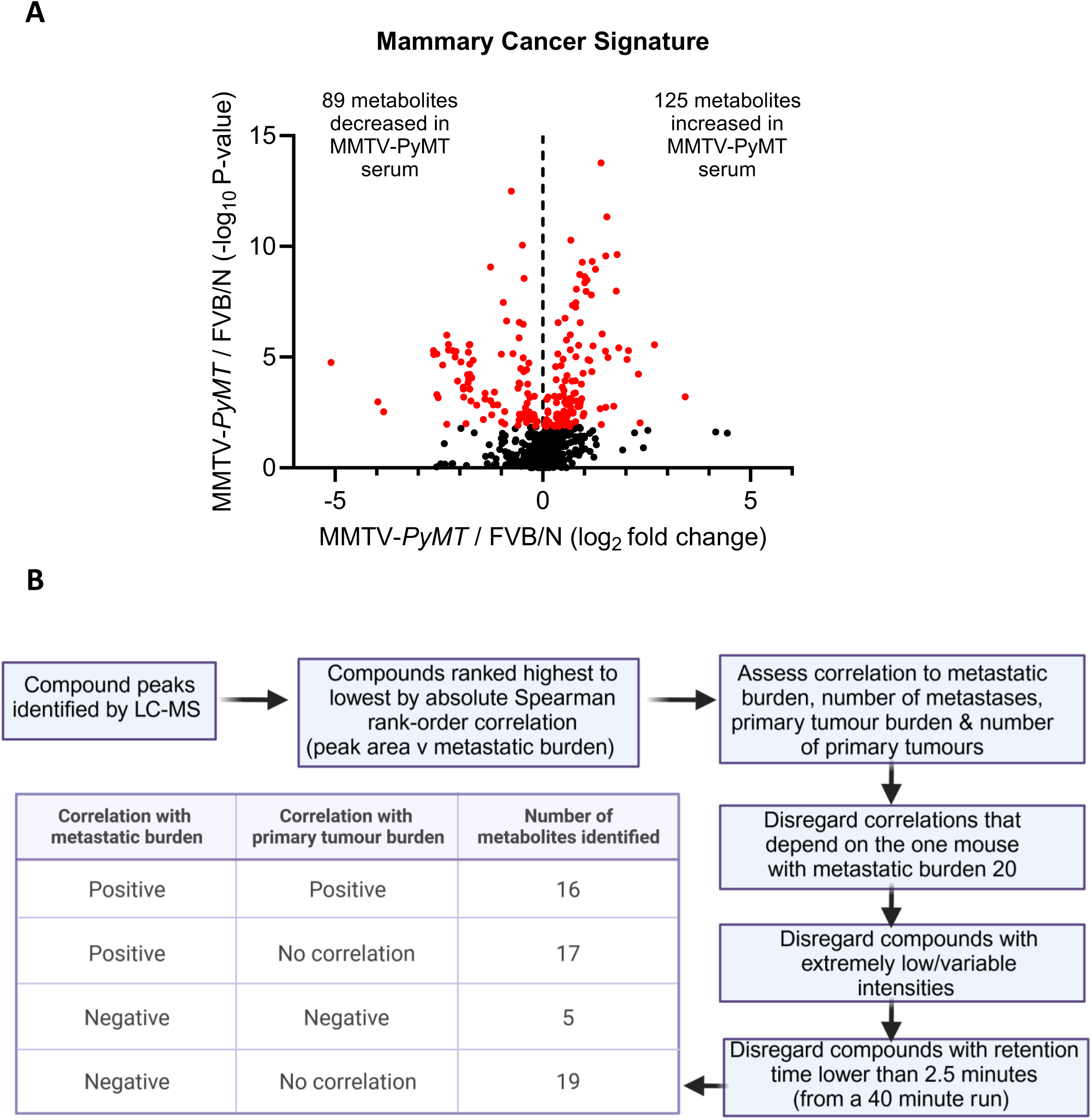
The circulating metabolic signature of mammary cancer. (A) Volcano plot representing compounds identified in the serum of FVB/N (n=9) and MMTV-*PyMT* tumour-bearing mice (n=29). From 740 compounds detected, 214 compounds (highlighted in red) were determined as being statistically significantly different in the serum of MMTV-*PyMT* mice compared to FVB/N WT, p-values calculated with unpaired t-test and adjusted with Benjamin Hochberg false discovery rate calculation. (B) Schematic representation of the methodology used to identify metabolites that correlate with metastasis independent of primary tumour burden. Molecular weights, retention times and compound identifications confirmed by comparison to commercial standard for each group are detailed in Table S1.

**Supplementary Figure 2:**
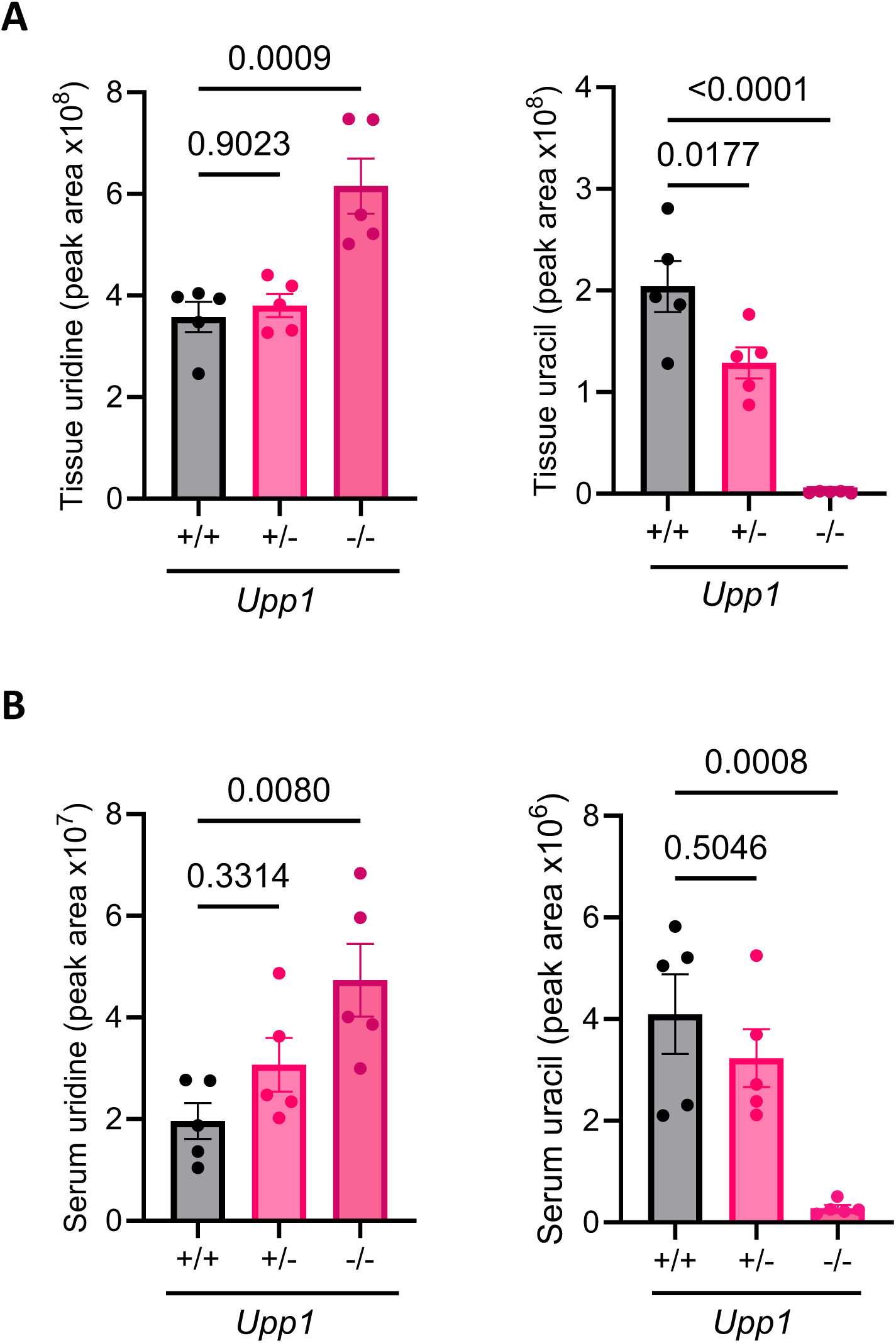
The metabolic characterisation of *Upp1* knockout mice. Polar metabolites were extracted from (A) tissue and (B) serum of FVB/N *Upp1^+/+^*, *Upp1^+/-^*, and *Upp1^-/-^* mice, and uridine and uracil levels were assessed by LC-MS. Tissue in this instance refer to pancreas. N=5 mice per experimental group, both sexes were used at 9 – 12 weeks of age, one-way ANOVA, mean ± SEM.

**Supplementary Figure 3:**
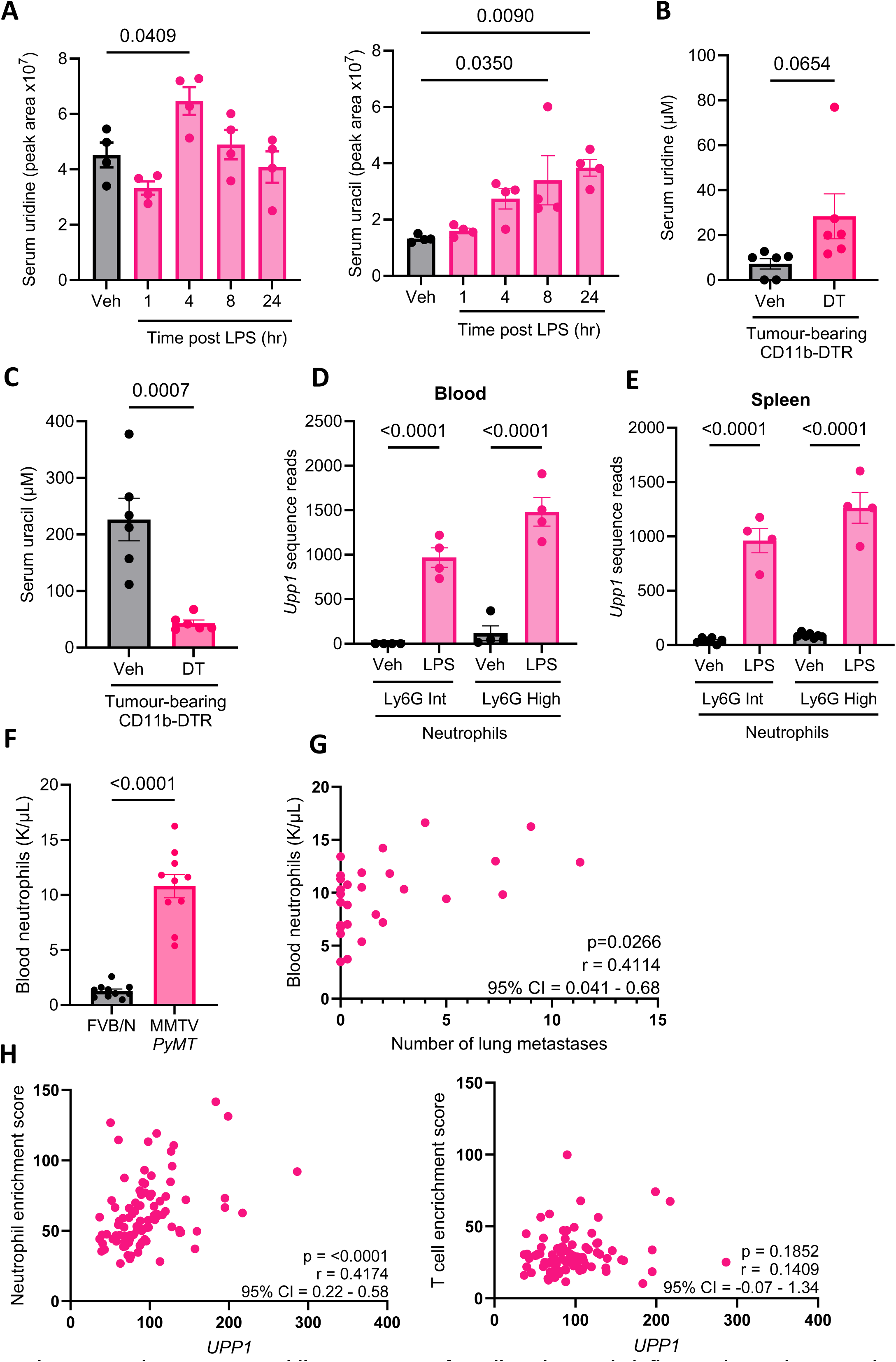
Neutrophils are a source of uracil, and *Upp1*, in inflammation and metastatic cancer. (A) FVB/N mice were dosed intraperitoneal (IP) with 0.5mg/kg lipopolysaccharide (LPS), or vehicle control, and serum uridine and uracil measured by LC-MS at defined timepoints post dosing, n=4 per experimental arm, one-way ANOVA, mean ± SEM. (B-C) Orthotopic PDAC tumours were established from 5×10^4^ KPC-7940B PDAC cells injected into the pancreas in 50µL of 1:1 Matrigel-DMEM + 10%FBS of F1 C57BL/6J CD11b-DTR mice and allowed to establish tumours for 14 days. Mice were then treated with diphtheria toxin (DT) at a concentration of 25ng/g, or vehicle, intraperitoneal. Mice were humanely euthanised 24 hours after treatment and serum uridine and uracil were assessed by LC-MS. N=6 mice per experimental group, unpaired t-test, mean ± SEM. (D-E) *Upp1* transcript reads detected by RNA-Seq in CD11b^+^Ly6G^+^ neutrophils flow cytometry sorted from the blood and spleen of C57BL/6 mice 24 hours post IP dosing of 1mg/kg LPS. Cell surface levels of Ly6G were determined as intermediate (immature neutrophils) or high (mature neutrophils) by flow cytometry. One-way ANOVA, mean ± SEM, n=4-7 mice per experimental group, with individual mice indicated by dot, described in Mackey et al., 2021 [30]. (F) Number of blood neutrophils were determined by IDEXX in 14-week-old female MMTV-*PyMT* tumour bearing mice (n=10) and age-matched FVB/N controls (n=10), unpaired t-test, ± SEM. (G) Number of blood neutrophils determined by IDEXX in female MMTV-*PyMT* tumour bearing mice at clinical endpoint (n=29), compared to number of lung metastases determined by immunohistology analysis of 3 lung sections. Spearman correlations are presented in graph (p value, Spearman r and 95% confidence interval). (H) Neutrophil enrichment score, and T cell enrichment score, were determined using Microenvironment Cell Populations counter (MCP) methodology [31] on human colon cancer dataset GSE33113. Spearman correlations are presented in graph. In all cases each dot refers to an individual mouse or human as appropriate.

**Supplementary Figure 4:**
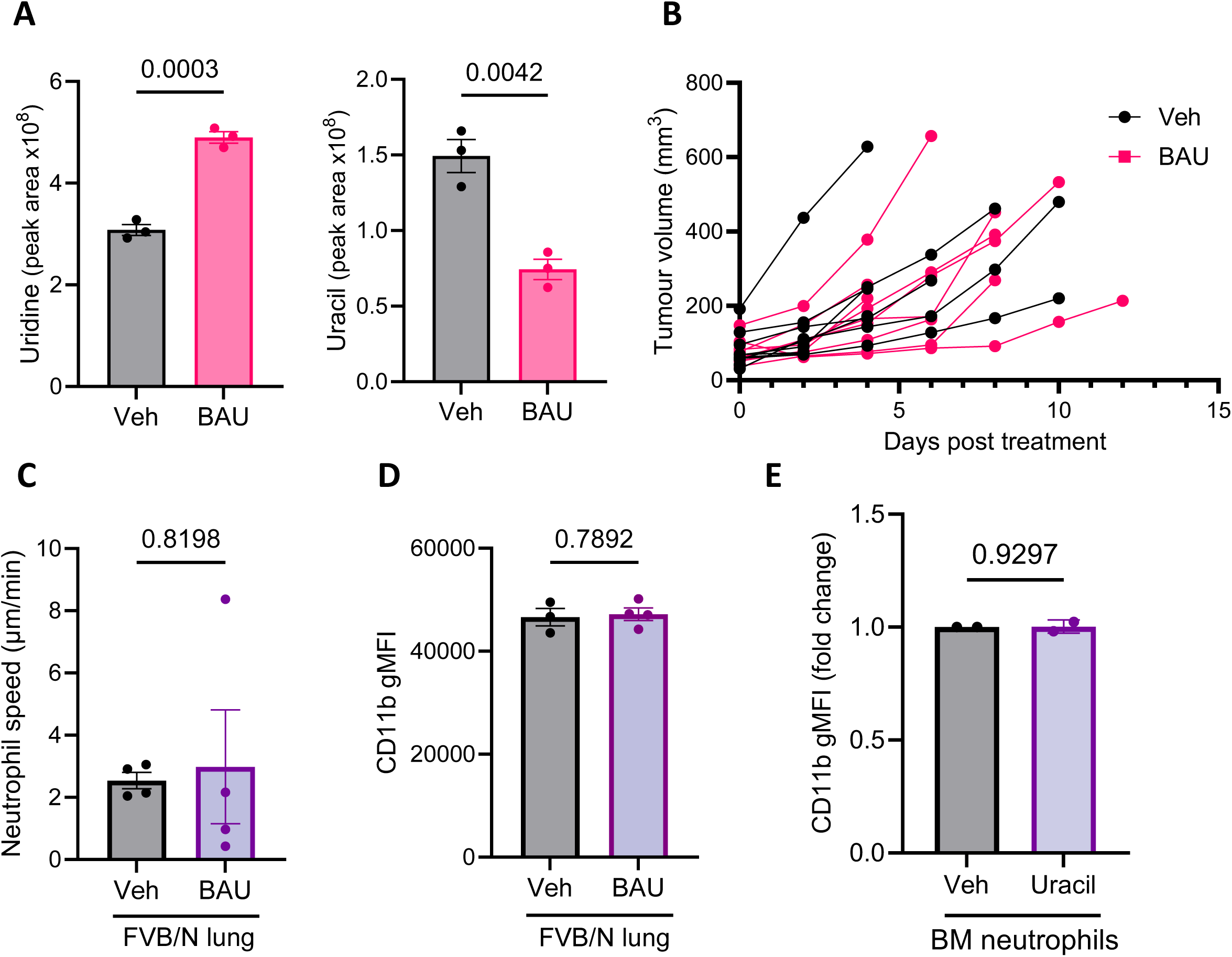
5-Benzylacyclouridine inhibits the metabolic activity of UPP1 but does not affect primary tumour growth. (A) *In vivo* validation of 5-benzylacyclouridine (BAU) as an inhibitor of UPP1. C57Bl/6 mice (n=3 per experimental group) were treated for 3 days with vehicle (0.5% HPMC/0.1% Tween:DMSO, 90:10; p.o., b.i.d) or BAU (30 mg/kg p.o., b.i.d.). Tissues were harvested, and uridine and uracil levels assessed by LC-MS. Tissue presented is from pancreas, unpaired t-test, mean ± SEM. (B) Tumour volumes calculated from female mice transplanted orthotopically with tumour fragments obtained from KP mice. Mice were dosed with vehicle (n=10) or BAU (n=9) following detection of measurable tumours. Tumour measurements were determined 3 times per week by calliper measurement. (C) Neutrophil speed was assessed in the lungs of FVB/N mice by live cell imaging of precision cut lung slices, comparing mice treated with vehicle or BAU for 3 consecutive days with mice harvested on the 4^th^ day, n=4 per experimental group, unpaired t-test, mean ± SEM. (D) Cell surface levels of CD11b on neutrophils (CD45^+^Ly6G^+^CD11b^+^) from the lungs of mice described in C, but with lung neutrophils assessed by flow cytometry, n=3-4 per experimental group, unpaired t-test, mean ± SEM. (E) Neutrophils were isolated from the bone marrow (BM) of mice, treated with 30µM uracil for 24 hours at 37°C/5%CO_2_ and then cell surface CD11b levels were assessed by flow cytometry, preparations from n=2 experimental mice, mean ± Std Dev, paired t-test. In each case, dots within bar graphs represent an individual mouse.

**Supplementary Figure 5:**
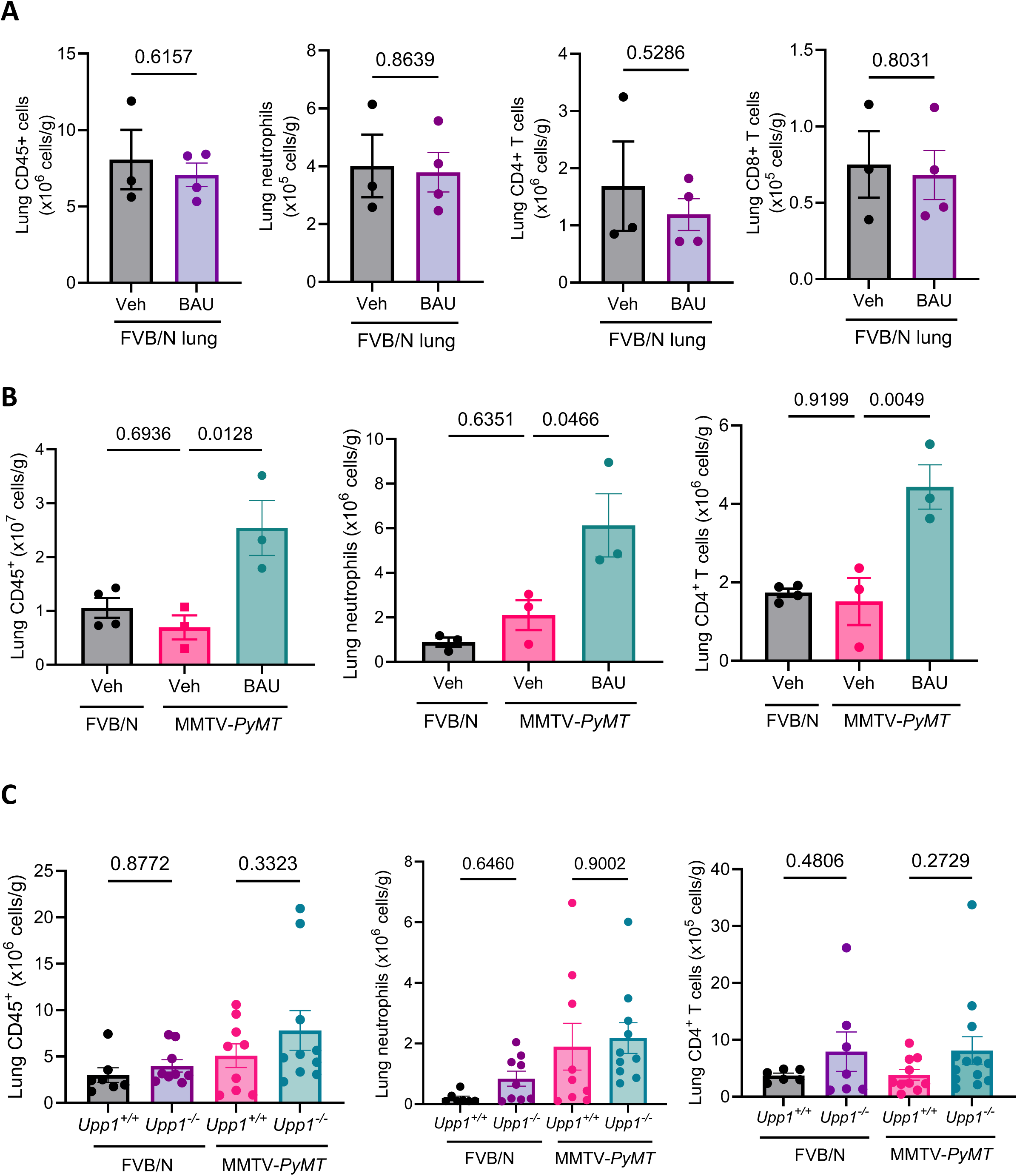
The effect of UPP1 on the immune landscape of the lung. (A) Cell number per gram of lung for CD45^+^ cells, neutrophils, CD4^+^ T cells and CD8^+^ T cells in FVB/N mice treated with vehicle (0.5% HPMC/0.1% Tween:DMSO, 90:10; p.o., b.i.d) or BAU (30 mg/kg p.o., b.i.d.) for 3 consecutive days with mice harvested on the 4^th^ day, assessed by flow cytometry, n=3-4 per group denoted by dots on graph, unpaired t-test, mean ± SEM. (B) Cell number per gram of lung for CD45^+^, neutrophils (Ly6G^+^CD11b^+^), and CD4^+^ T cells, assessed by flow cytometry, in the lungs of MMTV-*PyMT* mice treated with vehicle or BAU from palpable tumour until mammary tumours reached 10 – 15mm in diameter, including FVB/N mice treated with vehicle for time-matched periods, n=3–4 per experimental group denoted by dots on graph, one-way ANOVA, mean ± SEM. (C) Cell number per gram of lung for CD45^+^, neutrophils (Ly6G^+^CD11b^+^), and CD4^+^ T cells, assessed by flow cytometry, in the lungs of *Upp1^+/+^* and *Upp1^-/-^* mice, with MMTV-*PyMT* mice bearing mammary tumours 10 – 15mm in diameter, n=6–13 per experimental group denoted by dots on graph, one-way ANOVA, mean ± SEM.

**Supplementary Figure 6:**
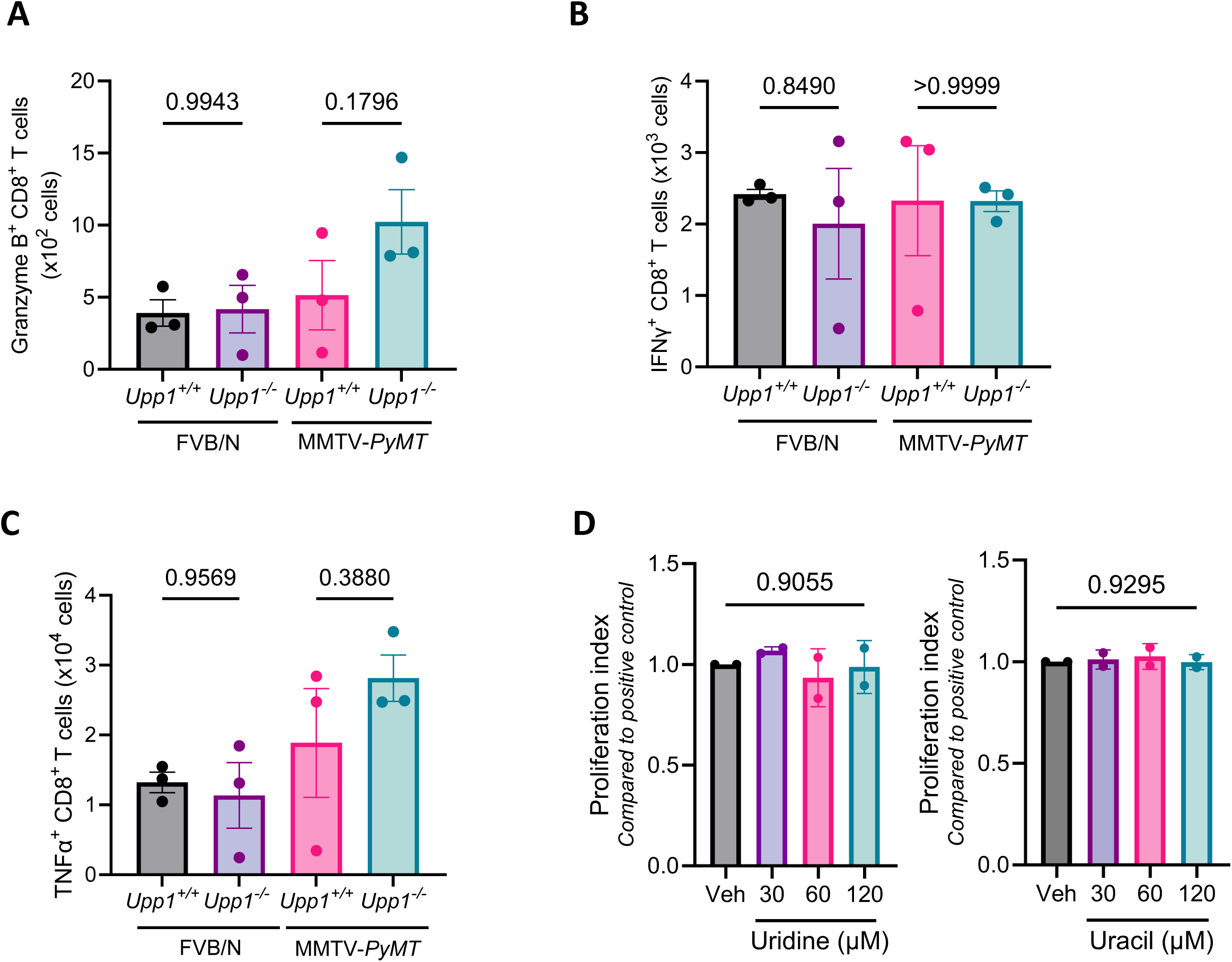
Understanding T cell effector function in the absence of *Upp1*. (A) Lungs of MMTV-*PyMT*;*Upp1^+/+^* and MMTV-*PyMT*;*Upp1^-/-^* tumour-bearing mice were harvested when one tumour measured 10–15mm diameter. Cells were prepared for flow cytometry, but with a 3 hour stimulation with PMA and ionomycin together with Brefeldin A when necessary, to enable assessment of intracellular Granzyme B levels, n=3 mice per experimental group, one-way ANOVA, mean ± SEM. (B) As described in A, but with assessment of intracellular Interferon-γ levels, one-way ANOVA, mean ± SEM. (C) As described in A, but with assessment of intracellular TNF-α levels, one-way ANOVA, mean ± SEM. (D) T cells were isolated from FVB/N mice, incubated in culture with CD3/CD28 dynabeads to stimulate proliferation in medium containing 30, 60 and 120µM uridine or uracil. Proliferation index was calculated compared to the positive control of T cells and vehicle alone stimulated with beads, cell preparations from n=2 biological repeats, mean ± Std Dev, unpaired t-test.

**Supplementary Figure 7.**
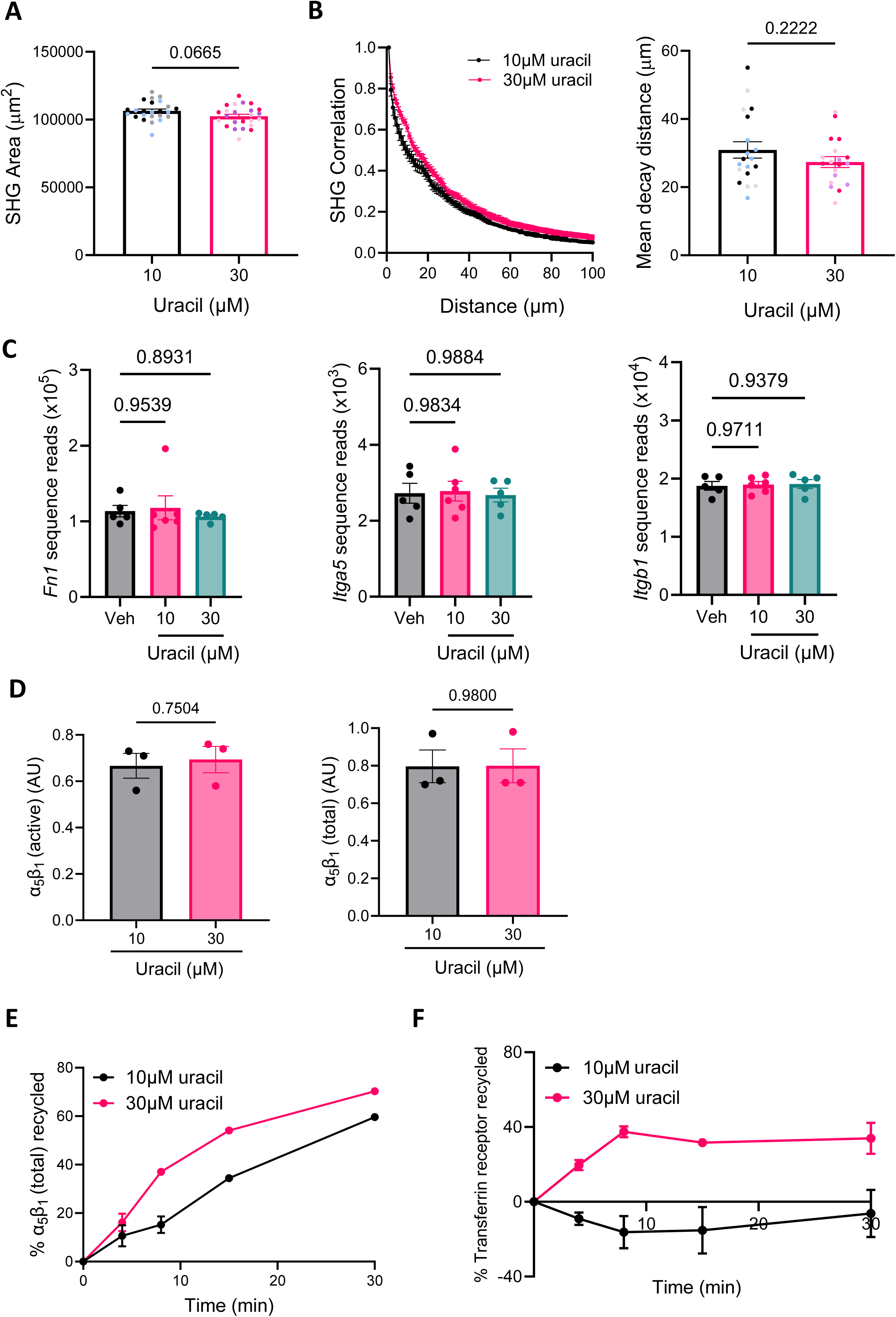
Understanding the extent to which uracil can influence fibroblast deposited extracellular matrix. (A) Fibrillar collagen was imaged by second harmonic generation (SHG) microscopy in organotypic plugs of rat tail collagen contracted by fibroblasts supplemented with 10 or 30µM uracil. A threshold was applied to the SHG signal and the area of SHG coverage per field of view was determined. The organisation of fibrillar collagen in each field of view was assessed by applying grey level co-occurrence matrix (GLCM), a second-order statistical method, described in further detail in [27]. (B) The mean correlation decay curves from each experimental condition, and the mean of the decay distances derived from those curves are presented, n = 24 fields of view across n=3 biological experimental repeats, with data from each experiment colour coded accordingly, mean ± SEM, unpaired t-test. N=19 fields of view are represented for mean decay distance as 6 fields of view did not behave like 2 phase decay curves and so could not be fitted in the equation. (C) RNA-Sequencing transcript reads from fibroblasts treated with vehicle, 10µM and 30µM uracil for 24 hours, for fibronectin (*Fn1*), integrin α_5_ (*Itga5*) and integrin β_1_ (*Itgb1*), n=5-6 biological repeats, one-way ANOVA, mean ± SEM. (D) Total protein levels quantified from total cell lysate via ELISA, arbitrary units (AU), n=3 biological repeats, unpaired t-test, mean ± SEM. (E) Recycling of the total levels of the fibronectin receptor, α_5_β_1_ integrin, assessed in fibroblasts treated with 10µM or 30µM uracil for 24 hours, data presented are the mean ± SEM from n=3 biological repeats. (F) Recycling of transferrin receptor assessed in fibroblasts treated with 10µM or 30µM uracil for 24 hours, data presented are the mean ± SEM from n=3 biological repeats.

**Supplementary Figure 8:**
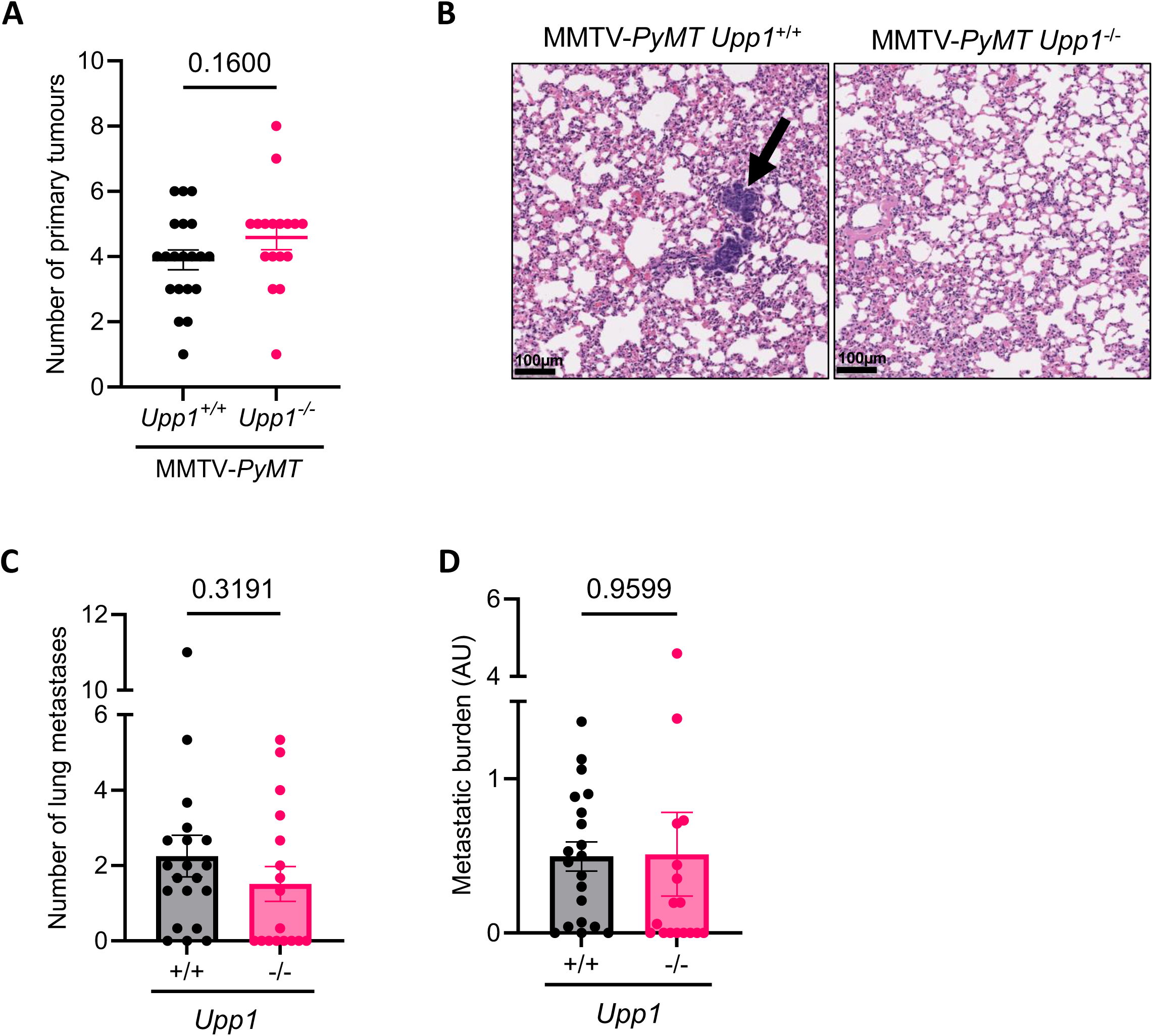
UPP1 does not influence primary tumour growth, but does influence metastasis, in the MMTV-*PyMT* model of mammary cancer. (A) Total number of primary mammary tumours at clinical endpoint in MMTV-*PyMT*;*Upp1^+/+^* and MMTV-*PyMT*;*Upp1^-/-^* mice, n= 17-20 mice per experimental condition (denoted by dots on graph for figures A, C and D), unpaired t-test, mean ±SEM. (B) Representative H&E stains for the lungs of MMTV-*PyMT*;*Upp1^+/+^* and MMTV-*PyMT*;*Upp1^-/-^* mice at clinical endpoint, scale bar represents 100µm. Arrow highlights a metastatic lesion. (C) Average number of metastases across 3 H&E sections per mouse, for MMTV-*PyMT*;*Upp1^+/+^*and MMTV-*PyMT*;*Upp1^-/-^* mice, unpaired t-test, mean ±SEM. (D) Average metastatic burden measured across 3 H&E sections per mouse, for MMTV-*PyMT*;*Upp1^+/+^* and MMTV-*PyMT*;*Upp1^-/-^* mice, unpaired t-test, mean ±SEM.

**Supplementary Table 1:** Serum metabolites that correlate with metastasis in the MMTV-*PyMT* model of mammary cancer. Spearman Correlation statistics are presented: Spearman r, 95% confidence interval, individual p value non-FDR corrected. Confirmed compound identifications are those that were validated against a commercial standard.

| Spearman rank correlation (r)<br>Compound Peak Area v<br>Metastatic Burden | 95% Confidence<br>interval | P Value | Correlation with<br>metastatic burden | Correlation with<br>primary tumour<br>burden | Molecular<br>Weight | Retention Time<br>(minutes) | Confirmed Compound<br>Identification |
| --- | --- | --- | --- | --- | --- | --- | --- |
| 0.6138 | 0.3088 to 0.8043 | 0.0004 | Positive | Positive | 225.07 | 6.97 |  |
| 0.5852 | 0.2679 to 0.7880 | 0.0009 | Positive | Positive | 145.07 | 10.50 |  |
| 0.5827 | 0.2644 to 0.7866 | 0.0009 | Positive | Positive | 201.11 | 8.24 |  |
| 0.5359 | 0.1999 to 0.7591 | 0.0027 | Positive | Positive | 285.10 | 5.50 | N4-Acetylcytidine |
| 0.5263 | 0.1870 to 0.7534 | 0.0034 | Positive | Positive | 114.04 | 8.10 |  |
| 0.5157 | 0.1729 to 0.7470 | 0.0042 | Positive | Positive | 153.05 | 5.50 |  |
| 0.5085 | 0.1635 to 0.7427 | 0.0049 | Positive | Positive | 515.29 | 2.93 |  |
| 0.4121 | 0.04241 to 0.6826 | 0.0263 | Positive | Positive | 132.05 | 9.05 | 3-Ureidopropionic acid |
| 0.4777 | 0.1236 to 0.7239 | 0.0088 | Positive | Positive | 172.01 | 13.07 | Glycerol 3-phosphate |
| 0.4769 | 0.1227 to 0.7234 | 0.0089 | Positive | Positive | 327.12 | 12.45 |  |
| 0.4644 | 0.1067 to 0.7156 | 0.0112 | Positive | Positive | 146.07 | 6.83 |  |
| 0.4429 | 0.07993 to 0.7022 | 0.0161 | Positive | Positive | 189.11 | 13.28 |  |
| 0.4200 | 0.05187 to 0.6876 | 0.0233 | Positive | Positive | 158.04 | 11.43 |  |
| 0.4185 | 0.05008 to 0.6867 | 0.0239 | Positive | Positive | 174.10 | 11.19 |  |
| 0.4173 | 0.04859 to 0.6859 | 0.0243 | Positive | Positive | 244.07 | 9.59 | Pseudouridine |
| 0.3951 | 0.02204 to 0.6715 | 0.0339 | Positive | Positive | 532.32 | 2.92 |  |
| 0.4254 | 0.05846 to 0.6911 | 0.0214 | Positive | No correlation | 208.05 | 9.61 |  |
| 0.5509 | 0.2203 to 0.7680 | 0.002 | Positive | No correlation | 352.08 | 13.05 |  |
| 0.5307 | 0.1929 to 0.7560 | 0.0031 | Positive | No correlation | 299.19 | 5.23 |  |
| 0.5001 | 0.1525 to 0.7376 | 0.0057 | Positive | No correlation | 210.04 | 12.21 |  |
| 0.4851 | 0.1331 to 0.7284 | 0.0077 | Positive | No correlation | 130.03 | 12.26 |  |
| 0.4841 | 0.1318 to 0.7278 | 0.0078 | Positive | No correlation | 266.05 | 9.65 |  |
| 0.4806 | 0.1274 to 0.7257 | 0.0083 | Positive | No correlation | 326.22 | 2.63 |  |
| 0.4651 | 0.1076 to 0.7161 | 0.011 | Positive | No correlation | 279.08 | 12.13 |  |
| 0.4604 | 0.1017 to 0.7132 | 0.012 | Positive | No correlation | 246.09 | 7.08 |  |
| 0.4584 | 0.09927 to 0.7119 | 0.0124 | Positive | No correlation | 113.05 | 3.27 |  |
| 0.4330 | 0.06769 to 0.6959 | 0.019 | Positive | No correlation | 131.07 | 12.70 |  |
| 0.4284 | 0.06206 to 0.6930 | 0.0204 | Positive | No correlation | 154.00 | 7.46 |  |
| 0.4178 | 0.04919 to 0.6862 | 0.0241 | Positive | No correlation | 112.03 | 5.73 | Uracil |
| 0.4160 | 0.04710 to 0.6851 | 0.0248 | Positive | No correlation | 148.04 | 12.23 |  |
| 0.4111 | 0.04116 to 0.6819 | 0.0267 | Positive | No correlation | 481.35 | 5.21 |  |
| 0.3983 | 0.02585 to 0.6736 | 0.0324 | Positive | No correlation | 129.04 | 5.93 |  |
| 0.3857 | 0.01099 to 0.6654 | 0.0388 | Positive | No correlation | 357.16 | 13.10 |  |
| -0.5522 | -0.7687 to -0.2220 | 0.0019 | Negative | Negative | 146.02 | 13.26 | Alpha-ketoglutarate |
| -0.4959 | -0.7351 to -0.1471 | 0.0062 | Negative | Negative | 466.31 | 2.55 |  |
| -0.5753 | -0.7823 to -0.2540 | 0.0011 | Negative | Negative | 779.55 | 2.76 |  |
| -0.5605 | -0.7737 to -0.2335 | 0.0016 | Negative | Negative | 519.33 | 5.18 |  |
| -0.5223 | -0.7510 to -0.1817 | 0.0037 | Negative | Negative | 102.03 | 13.27 |  |
| -0.5689 | -0.7786 to -0.2451 | 0.0013 | Negative | No correlation | 607.35 | 3.02 |  |
| -0.5605 | -0.7689 to -0.2223 | 0.0019 | Negative | No correlation | 541.32 | 3.03 |  |
| -0.5470 | -0.7657 to -0.2149 | 0.0021 | Negative | No correlation | 236.09 | 5.24 |  |
| -0.5298 | -0.7555 to -0.1918 | 0.0031 | Negative | No correlation | 134.02 | 14.18 |  |
| -0.5041 | -0.7400 to -0.1577 | 0.0053 | Negative | No correlation | 246.10 | 5.25 |  |
| -0.4905 | -0.7317 to -0.1401 | 0.0069 | Negative | No correlation | 548.18 | 12.67 |  |
| -0.4903 | -0.7316 to -0.1397 | 0.0069 | Negative | No correlation | 384.14 | 12.54 |  |
| -0.4612 | -0.7136 to -0.1027 | 0.0118 | Negative | No correlation | 304.14 | 13.50 |  |
| -0.4543 | -0.7093 to -0.09403 | 0.0133 | Negative | No correlation | 246.10 | 3.23 |  |
| -0.4543 | -0.7093 to -0.09403 | 0.0133 | Negative | No correlation | 868.57 | 2.77 |  |
| -0.4444 | -0.7031 to -0.08176 | 0.0157 | Negative | No correlation | 148.04 | 13.06 |  |
| -0.4424 | -0.7019 to -0.07932 | 0.0163 | Negative | No correlation | 545.35 | 5.17 |  |
| -0.4390 | -0.6997 to -0.07506 | 0.0172 | Negative | No correlation | 369.12 | 12.62 |  |
| -0.4318 | -0.6952 to -0.06628 | 0.0193 | Negative | No correlation | 567.33 | 5.13 |  |
| -0.4195 | -0.6873 to -0.05127 | 0.0235 | Negative | No correlation | 549.18 | 12.60 |  |
| -0.4079 | -0.6798 to -0.03732 | 0.0281 | Negative | No correlation | 275.15 | 15.19 |  |
| -0.4054 | -0.6783 to -0.03437 | 0.0291 | Negative | No correlation | 261.12 | 10.30 |  |
| -0.4032 | -0.6768 to -0.03172 | 0.0301 | Negative | No correlation | 132.04 | 12.48 |  |
| -0.3921 | -0.6696 to -0.01854 | 0.0354 | Negative | No correlation | 175.05 | 12.28 | N-Acetylaspartic acid |

**Supplementary Table 2:** Antibody panel for neutrophil characterisation.

| Target | Fluorophore | Dilution | Supplier | Catalogue Number |
| --- | --- | --- | --- | --- |
| **Zombie Green** | Zombie Green | 1/1000 | Biolegend | 423111 |
| **Ter119** | FITC (Dump) | 1/200 | Biolegend | 116206 |
| **Nkp46** | FITC (Dump) | 1/200 | Biolegend | 137606 |
| **CD115 (CSF1R)** | FITC (Dump) | 1/200 | TONBO bioscience | 35-1152-u100 |
| **CD3** | FITC (Dump) | 1/200 | Biolegend | 100306 |
| **CD19** | FITC (Dump) | 1/200 | Biolegend | 101506 |
| **CD45** | BV650 | 1/200 | Biolegend | 103151 |
| **CD11b** | BV605 | 1/200 | Biolegend | 101257 |
| **Ly6G** | Bv510 | 1/200 | Biolegend | 127633 |
| **CD54** | APC | 1/200 | Biolegend | 116120 |
| **CD62L** | BUV395 | 1/200 | Biolegend | 740218 |
| **CD101** | PE-Cy7 | 1/200 | Life Tech (Invitrogen) | 25-1011-82 |
| **CD117** | Bv711 | 1/100 | Biolegend | 105835 |
| **CXCR4** | BV421 | 1/100 | Biolegend | 146511 |
| **CXCR2** | PE | 1/50 | Biolegend | 149304 |
| **CD63** | Percp-Cy5.5 | 1/200 | Biolegend | 143912 |
| **Siglec-F** | APCCy7 | 1/200 | BD bioscience | 565527 |

**Supplementary Table 3:** Antibody panel for lung immunophenotyping.

| Target | Fluorophore | Dilution | Supplier | Catalogue Number |
| --- | --- | --- | --- | --- |
| **Zombie NIR** | Zombie NIR | 1/1000 | Biolegend | 423105 |
| **CD45** | AF700 | 1/200 | Biolegend | 103128 |
| **CD19** | Bv605 | 1/200 | Biolegend | 115540 |
| **CD3** | PercpCy5.5 | 1/200 | Biolegend | 100328 |
| **CD8** | FITC | 1/200 | Biolegend | 100706 |
| **CD4** | PeCy7 | 1/200 | Biolegend | 116016 |
| **gdTCR** | PE | 1/200 | Biolegend | 118108 |
| **SiglecF** | AF647 | 1/200 | BD Pharmigen | 562680 |
| **CD115** | Bv421 | 1/200 | Biolegend | 135513 |
| **CD11b** | Bv650 | 1/200 | Biolegend | 101259 |
| **Ly6G** | Bv711 | 1/200 | Biolegend | 127463 |
| **F4/80** | Bv510 | 1/200 | Biolegend | 123135 |

**Supplementary Table 4:** Antibody panel for lymphoid characterisation.

| Target | Fluorophore | Dilution | Supplier | Catalogue Number |
| --- | --- | --- | --- | --- |
| **zombie NIR** | Zombie NIR | 1/1000 | Biolegend | 423105 |
| **CD11b, CD11c, CD19** | BV605 (Dump) | 1/200 | Biolegend | 101257, 117334, 115540 |
| **PD-1** | APC | 1/200 | Biolegend | 135210 |
| **CD45** | A700 | 1/200 | Biolegend | 103128 |
| **gdTCR** | PE | 1/200 | Biolegend | 118108 |
| **CD62L** | PE-Dazzle | 1/200 | Biolegend | 104448 |
| **CD4** | PE-Cy7 | 1/200 | Biolegend | 116016 |
| **CD8** | PerCP-Cy5.5 | 1/200 | Biolegend | 100734 |
| **CD44** | BUV395 | 1/200 | BD Horizon | 568507 |
| NKp46 | BV421 | 1/200 | Biolegend | 137611 |
| CD27 | BV510 | 1/200 | Biolegend | 124229 |
| CD69 | BV650 | 1/200 | Biolegend | 104541 |
| ICOS | BV711 | 1/200 | Biolegend | 313548 |
| CD3 | BV785 | 1/200 | Biolegend | 100355 |

**Supplementary Table 5:**
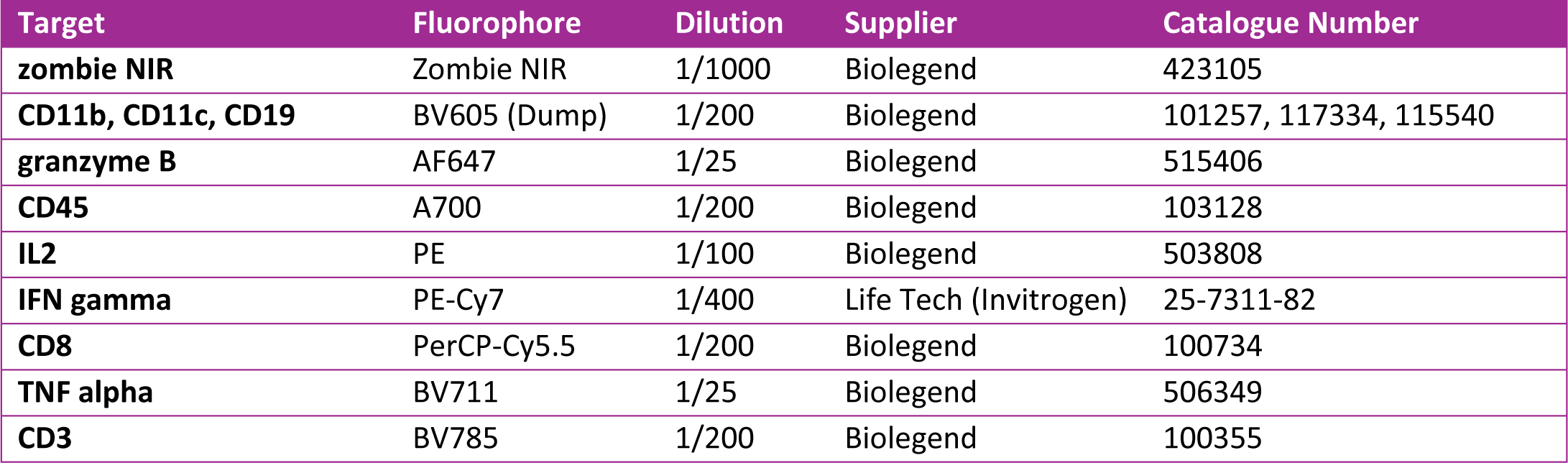
Antibody panel for intracellular characterisation of T cell effector function.

| Target | Fluorophore | Dilution | Supplier | Catalogue Number |
| --- | --- | --- | --- | --- |
| zombie NIR | Zombie NIR | 1/1000 | Biolegend | 423105 |
| CD11b, CD11c, CD19 | BV605 (Dump) | 1/200 | Biolegend | 101257, 117334, 115540 |
| granzyme B | AF647 | 1/25 | Biolegend | 515406 |
| CD45 | A700 | 1/200 | Biolegend | 103128 |
| IL2 | PE | 1/100 | Biolegend | 503808 |
| IFN gamma | PE-Cy7 | 1/400 | Life Tech (Invitrogen) | 25-7311-82 |
| CD8 | PerCP-Cy5.5 | 1/200 | Biolegend | 100734 |
| TNF alpha | BV711 | 1/25 | Biolegend | 506349 |
| CD3 | BV785 | 1/200 | Biolegend | 100355 |

**Supplementary Table 6:**
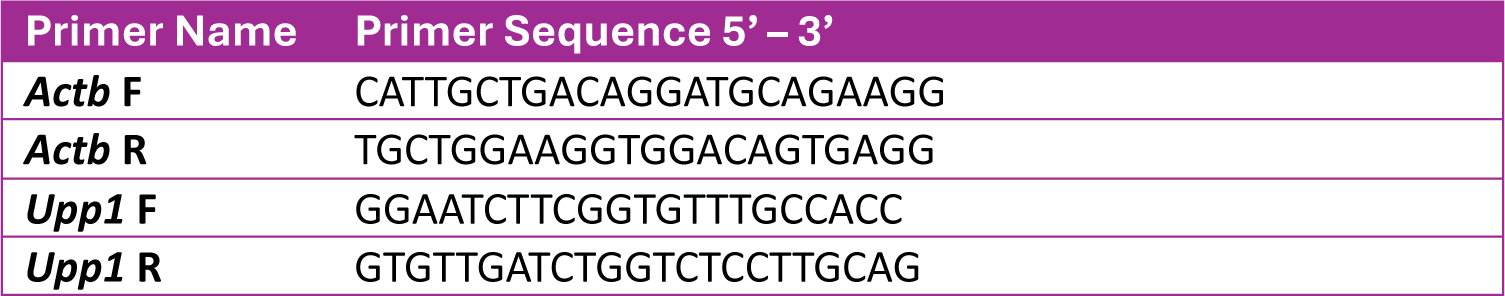
qRT-PCR primer sequences.

**Supplementary Table 7:** Isotopically labelled internal standards for CD11b-DTR metabolomics.

| Standard | Supplier information |
| --- | --- |
| <sup>13</sup>C-labeled yeast extract | (Cambridge Isotope Laboratory, Andover, MA, ISO1) |
| <sup>2</sup>H<sub>9</sub> choline | (Cambridge Isotope Laboratory, Andover, MA, DLM-549) |
| <sup>13</sup>C<sub>4</sub> 3-hydroxybutyrate | (Cambridge Isotope Laboratory, Andover, MA, CLM-3853) |
| <sup>13</sup>C<sub>6</sub><sup>15</sup>N<sub>2</sub> cystine | (Cambridge Isotope Laboratory, Andover, MA, CNLM4244) |
| <sup>13</sup>C<sub>3</sub> lactate | (Sigma-Aldrich, Darmstadt, Germany, 485926) |
| <sup>13</sup>C<sub>6</sub> glucose | (Cambridge Isotope Laboratory, Andover, MA, CLM-1396) |
| <sup>13</sup>C<sub>3</sub> serine | (Cambridge Isotope Laboratory, Andover, MA, CLM-1574) |
| <sup>13</sup>C<sub>2</sub> glycine | (Cambridge Isotope Laboratory, Andover, MA, CLM-1017) |
| <sup>13</sup>C<sub>5</sub> hypoxanthine | (Cambridge Isotope Laboratory, Andover, MA, CLM8042) |
| <sup>13</sup>C<sub>2</sub><sup>15</sup>N taurine | (Cambridge Isotope Laboratory, Andover, MA, CNLM-10253) |
| <sup>13</sup>C<sub>3</sub> glycerol | (Cambridge Isotope Laboratory, Andover, MA, CLM-1510) |
| <sup>2</sup>H<sub>3</sub> creatinine | (Cambridge Isotope Laboratory, Andover, MA, DLM-3653) |

## Notes

### Competing Interest Statement

The authors have declared no competing interest.

## References

1. Chaffer, C.L. and R.A. Weinberg, A perspective on cancer cell metastasis. Science, 2011. 331(6024): p. 1559–64.

2. Gupta, G.P. and J. Massague, Cancer metastasis: building a framework. Cell, 2006. 127(4): p. 679–95.

3. Riggio, A.I., K.E. Varley, and A.L. Welm, The lingering mysteries of metastatic recurrence in breast cancer. Br J Cancer, 2021. 124(1): p. 13–26.

4. Welch, D.R. and D.R. Hurst, Defining the Hallmarks of Metastasis. Cancer Res, 2019. 79(12): p. 3011–3027.

5. Shojaei, F., et al., Role of Bv8 in neutrophil-dependent angiogenesis in a transgenic model of cancer progression. Proc Natl Acad Sci U S A, 2008. 105(7): p. 2640–5.

6. Shojaei, F., et al., Bv8 regulates myeloid-cell-dependent tumour angiogenesis. Nature, 2007. 450(7171): p. 825–31.

7. Coffelt, S.B., et al., IL-17-producing gammadelta T cells and neutrophils conspire to promote breast cancer metastasis. Nature, 2015. 522(7556): p. 345–348.

8. Wculek, S.K. and I. Malanchi, Neutrophils support lung colonization of metastasis-initiating breast cancer cells. Nature, 2015. 528(7582): p. 413–7.

9. Casbon, A.J., et al., Invasive breast cancer reprograms early myeloid differentiation in the bone marrow to generate immunosuppressive neutrophils. Proc Natl Acad Sci U S A, 2015. 112(6): p. E566–75.

10. Jackstadt, R., et al., Epithelial NOTCH Signaling Rewires the Tumor Microenvironment of Colorectal Cancer to Drive Poor-Prognosis Subtypes and Metastasis. Cancer Cell, 2019. 36(3): p. 319–336 e7.

11. Steele, C.W., et al., CXCR2 Inhibition Profoundly Suppresses Metastases and Augments Immunotherapy in Pancreatic Ductal Adenocarcinoma. Cancer Cell, 2016. 29(6): p. 832–845.

12. Li, P., et al., Dual roles of neutrophils in metastatic colonization are governed by the host NK cell status. Nat Commun, 2020. 11(1): p. 4387.

13. Hirai, H., et al., CCR1-mediated accumulation of myeloid cells in the liver microenvironment promoting mouse colon cancer metastasis. Clin Exp Metastasis, 2014. 31(8): p. 977–89.

14. Hoye, A.M. and J.T. Erler, Structural ECM components in the premetastatic and metastatic niche. Am J Physiol Cell Physiol, 2016. 310(11): p. C955–67.

15. Davis, R.T., et al., Transcriptional diversity and bioenergetic shift in human breast cancer metastasis revealed by single-cell RNA sequencing. Nat Cell Biol, 2020. 22(3): p. 310–320.

16. Gong, Z., et al., Lipid-laden lung mesenchymal cells foster breast cancer metastasis via metabolic reprogramming of tumor cells and natural killer cells. Cell Metab, 2022. 34(12): p. 1960–1976 e9.

17. Jeon, J.H., et al., Current Understanding on the Metabolism of Neutrophils. Immune Netw, 2020. 20(6): p. e46.

18. Rice, C.M., et al., Tumour-elicited neutrophils engage mitochondrial metabolism to circumvent nutrient limitations and maintain immune suppression. Nat Commun, 2018. 9(1): p. 5099.

19. Barnes, T.A. and E. Amir, HYPE or HOPE: the prognostic value of infiltrating immune cells in cancer. Br J Cancer, 2017. 117(4): p. 451–460.

20. Riaz, M., et al., High TWIST1 mRNA expression is associated with poor prognosis in lymph node-negative and estrogen receptor-positive human breast cancer and is co-expressed with stromal as well as ECM related genes. Breast Cancer Res, 2012. 14(5): p. R123.

21. DeBerardinis, R.J., et al., The biology of cancer: metabolic reprogramming fuels cell growth and proliferation. Cell Metab, 2008. 7(1): p. 11–20.

22. Guy, C.T., R.D. Cardiff, and W.J. Muller, Induction of mammary tumors by expression of polyomavirus middle T oncogene: a transgenic mouse model for metastatic disease. Mol Cell Biol, 1992. 12(3): p. 954–61.

23. Lin, E.Y., et al., Progression to malignancy in the polyoma middle T oncoprotein mouse breast cancer model provides a reliable model for human diseases. Am J Pathol, 2003. 163(5): p. 2113–26.

24. Attalla, S., et al., Insights from transgenic mouse models of PyMT-induced breast cancer: recapitulating human breast cancer progression in vivo. Oncogene, 2021. 40(3): p. 475–491.

25. Mei, Y., et al., Postponing tumor onset and tumor progression can be achieved by alteration of local tumor immunity. Cancer Cell Int, 2021. 21(1): p. 97.

26. Morton, J.P., et al., Mutant p53 drives metastasis and overcomes growth arrest/senescence in pancreatic cancer. Proc Natl Acad Sci U S A, 2010. 107(1): p. 246–51.

27. Novo, D., et al., Mutant p53s generate pro-invasive niches by influencing exosome podocalyxin levels. Nat Commun, 2018. 9(1): p. 5069.

28. Gyorffy, B., Transcriptome-level discovery of survival-associated biomarkers and therapy targets in non-small-cell lung cancer. Br J Pharmacol, 2024. 181(3): p. 362–374.

29. Gyorffy, B., Integrated analysis of public datasets for the discovery and validation of survival-associated genes in solid tumors. Innovation (Camb), 2024. 5(3): p. 100625.

30. Mackey, J.B.G., et al., Maturation, developmental site, and pathology dictate murine neutrophil function. bioRxiv, 2021: p. 2021.07.21.453108.

31. Becht, E., et al., Estimating the population abundance of tissue-infiltrating immune and stromal cell populations using gene expression. Genome Biol, 2016. 17(1): p. 218.

32. Fercoq, F., et al., Integrin inactivation slows down neutrophils congesting the pre-metastatic lung in a model of breast cancer. bioRxiv, 2024: p. 2024.03.19.585724.

33. Aarts, C.E.M., et al., Neutrophils as Suppressors of T Cell Proliferation: Does Age Matter? Front Immunol, 2019. 10: p. 2144.

34. Gillis, S., et al., Long-term culture of human antigen-specific cytotoxic T-cell lines. J Exp Med, 1978. 148(4): p. 1093–8.

35. Gillis, S. and K.A. Smith, Long term culture of tumour-specific cytotoxic T cells. Nature, 1977. 268(5616): p. 154–6.

36. Morgan, D.A., F.W. Ruscetti, and R. Gallo, Selective in vitro growth of T lymphocytes from normal human bone marrows. Science, 1976. 193(4257): p. 1007–8.

37. Smith, K.A., Interleukin-2: inception, impact, and implications. Science, 1988. 240(4856): p. 1169–76.

38. Zhang, C., et al., Fibrotic microenvironment promotes the metastatic seeding of tumor cells via activating the fibronectin 1/secreted phosphoprotein 1-integrin signaling. Oncotarget, 2016. 7(29): p. 45702–45714.

39. He, X.Y., et al., Chronic stress increases metastasis via neutrophil-mediated changes to the microenvironment. Cancer Cell, 2024. 42(3): p. 474–486 e12.

40. Dong, Q., et al., Pre-metastatic Niche Formation in Different Organs Induced by Tumor Extracellular Vesicles. Front Cell Dev Biol, 2021. 9: p. 733627.

41. Caswell, P.T., et al., Rab25 associates with alpha5beta1 integrin to promote invasive migration in 3D microenvironments. Dev Cell, 2007. 13(4): p. 496–510.

42. Sundararaman, A. and H. Mellor, A functional antagonism between RhoJ and Cdc42 regulates fibronectin remodelling during angiogenesis. Small GTPases, 2021. 12(4): p. 241–245.

43. Kaplan, R.N., et al., VEGFR1-positive haematopoietic bone marrow progenitors initiate the pre-metastatic niche. Nature, 2005. 438(7069): p. 820–7.

44. Tennant, D.A., R.V. Duran, and E. Gottlieb, Targeting metabolic transformation for cancer therapy. Nat Rev Cancer, 2010. 10(4): p. 267–77.

45. Nwosu, Z.C., et al., Uridine-derived ribose fuels glucose-restricted pancreatic cancer. Nature, 2023. 618(7963): p. 151–158.

46. Skinner, O.S., et al., Salvage of ribose from uridine or RNA supports glycolysis in nutrient-limited conditions. Nat Metab, 2023. 5(5): p. 765–776.

47. Li, Y., et al., UPP1 promotes lung adenocarcinoma progression through the induction of an immunosuppressive microenvironment. Nat Commun, 2024. 15(1): p. 1200.

48. Gross, E.T., et al., Immunosurveillance and immunoediting in MMTV-PyMT-induced mammary oncogenesis. Oncoimmunology, 2017. 6(2): p. e1268310.

49. Ajina, R., et al., Abstract 4584: Malignant pancreatic cancer cells respond to immune selection pressure to foster immunosuppression. Cancer Research, 2019. 79(13_Supplement): p. 4584–4584.

50. Yipp, B.G., et al., The Lung is a Host Defense Niche for Immediate Neutrophil-Mediated Vascular Protection. Sci Immunol, 2017. 2(10).

51. DiMilla, P.A., K. Barbee, and D.A. Lauffenburger, Mathematical model for the effects of adhesion and mechanics on cell migration speed. Biophys J, 1991. 60(1): p. 15–37.

52. Lishko, V.K., V.P. Yakubenko, and T.P. Ugarova, The interplay between integrins alphaMbeta2 and alpha5beta1 during cell migration to fibronectin. Exp Cell Res, 2003. 283(1): p. 116–26.

53. Paul, N.R., G. Jacquemet, and P.T. Caswell, Endocytic Trafficking of Integrins in Cell Migration. Curr Biol, 2015. 25(22): p. R1092–105.

54. Rossi, M., et al., PHGDH heterogeneity potentiates cancer cell dissemination and metastasis. Nature, 2022. 605(7911): p. 747-753.

55. Kelm, M., et al., Regulation of neutrophil function by selective targeting of glycan epitopes expressed on the integrin CD11b/CD18. FASEB J, 2020. 34(2): p. 2326–2343.

56. Gollapudi, S., et al., Steric pressure between glycosylated transmembrane proteins inhibits internalization by endocytosis. Proc Natl Acad Sci U S A, 2023. 120(15): p. e2215815120.

57. Fan, J., et al., Analysis of signature genes and association with immune cells infiltration in pediatric septic shock. Front Immunol, 2022. 13: p. 1056750.

58. Ross, S.H. and D.A. Cantrell, Signaling and Function of Interleukin-2 in T Lymphocytes. Annu Rev Immunol, 2018. 36: p. 411–433.

59. Li, P., et al., Lung mesenchymal cells elicit lipid storage in neutrophils that fuel breast cancer lung metastasis. Nat Immunol, 2020. 21(11): p. 1444–1455.

60. Paavola, K.J., et al., The Fibronectin-ILT3 Interaction Functions as a Stromal Checkpoint that Suppresses Myeloid Cells. Cancer Immunol Res, 2021. 9(11): p. 1283–1297.

61. Goncalves da Silva, E.F., et al., Therapeutic effect of uridine phosphorylase 1 (UPP1) inhibitor on liver fibrosis in vitro and in vivo. Eur J Pharmacol, 2021. 890: p. 173670.

62. Liu, X., et al., Somatic loss of BRCA1 and p53 in mice induces mammary tumors with features of human BRCA1-mutated basal-like breast cancer. Proc Natl Acad Sci U S A, 2007. 104(29): p. 12111–6.

63. Millar, R., et al., The MSP-RON axis stimulates cancer cell growth in models of triple negative breast cancer. Mol Oncol, 2020. 14(8): p. 1868–1880.

64. Villar, V.H., et al., Hepatic glutamine synthetase controls N(5)-methylglutamine in homeostasis and cancer. Nat Chem Biol, 2023. 19(3): p. 292–300.

65. Adams, K.J., et al., Skyline for Small Molecules: A Unifying Software Package for Quantitative Metabolomics. J Proteome Res, 2020. 19(4): p. 1447–1458.

66. Vande Voorde, J., et al., Improving the metabolic fidelity of cancer models with a physiological cell culture medium. Sci Adv, 2019. 5(1): p. eaau7314.

67. Sullivan, M.R., et al., Quantification of microenvironmental metabolites in murine cancers reveals determinants of tumor nutrient availability. Elife, 2019. 8.

68. Sullivan, M.R., C.A. Lewis, and A. Muir, Isolation and Quantification of Metabolite Levels in Murine Tumor Interstitial Fluid by LC/MS. Bio Protoc, 2019. 9(22): p. e3427.

69. Curio, S. and G.T. Belz, The unique role of innate lymphoid cells in cancer and the hepatic microenvironment. Cell Mol Immunol, 2022. 19(9): p. 1012–1029.

70. Timpson, P., et al., Organotypic collagen I assay: a malleable platform to assess cell behaviour in a 3-dimensional context. J Vis Exp, 2011(56): p. e3089.

71. Alshetaiwi, H., et al., Defining the emergence of myeloid-derived suppressor cells in breast cancer using single-cell transcriptomics. Sci Immunol, 2020. 5(44).

